# Fin whale (*Balaenoptera physalus*) mitogenomics: A cautionary tale of defining sub-species from mitochondrial sequence monophyly

**DOI:** 10.1101/488130

**Authors:** Andrea A. Cabrera, Jeroen P. A. Hoekendijk, Alex Aguilar, Susan G. Barco, Simon Berrow, Dorete Bloch, Asunción Borrell, Haydée A. Cunha, Luciano Dalla Rosa, Carolina P. Dias, Pauline Gauffier, Wensi Hao, Scott Landry, Finn Larsen, Vidal Martín, Sally Mizroch, Tom Oosting, Nils Øien, Christophe Pampoulie, Simone Panigada, Rui Prieto, Christian Ramp, Vania E. Rivera-Léon, Jooke Robbins, Conor Ryan, Elena Schall, Richard Sears, Mónica A. Silva, Jorge Urbán, Frederick W. Wenzel, Per J. Palsbøll, Martine Bérubé

**Author notes:** Corresponding author (A.A. Cabrera). Equally contributing authors (Andrea A. Cabrera, Jeroen P. A. Hoekendijk). senior authors (Per J. Palsbøll, Martine Bérubé). deceased (Dorete Bloch).

## Abstract

**Highlights:**

- Mitochondrial monophyly is commonly employed to define evolutionary significant units.
- Monophyly may be caused by insufficient sampling or a recent common ancestor.
- Mitogenomic studies are generally based on few samples and prone to sampling issues.
- Expanded mitogenome sampling negates previous monophyly in fin whales.

**Abstract:** The advent of massive parallel sequencing technologies has resulted in an increase of studies based upon complete mitochondrial genome DNA sequences that revisit the taxonomic status within and among species. Spatially distinct monophyly in mitogenomic genealogies, i.e., the sharing of a recent common ancestor among con-specific samples collected in the same region has been viewed as evidence for subspecies. Several recent studies in cetaceans have employed this criterion to suggest subsequent intraspecific taxonomic revisions. We reason that employing intra-specific, spatially distinct monophyly at non-recombining, clonally inherited genomes is an unsatisfactory criterion for defining subspecies based upon theoretical (genetic drift) and practical (sampling effort) arguments. This point is illustrated by a re-analysis of a global mitogenomic assessment of fin whales, *Balaenoptera physalus* spp., published by Archer et al. (2013) which proposed to further subdivide the Northern Hemisphere fin whale subspecies, *B. p. physalus*. The proposed revision was based upon the detection of spatially distinct monophyly among North Atlantic and North Pacific fin whales in a genealogy based upon complete mitochondrial genome DNA sequences. The extended analysis conducted in this study (1,676 mitochondrial control region, 162 complete mitochondrial genome DNA sequences and 20 microsatellite loci genotyped in 358 samples) revealed that the apparent monophyly among North Atlantic fin whales reported by Archer et al. (2013) to be due to low sample sizes. In conclusion, defining sub-species from monophyly (i.e., the absence of para-or polyphyly) can lead to erroneous conclusions due to relatively “trivial” aspects, such as sampling. Basic population genetic processes (i.e., genetic drift and migration) also affect the time to most recent common ancestor and hence the probability that individuals in a sample are monophyletic.

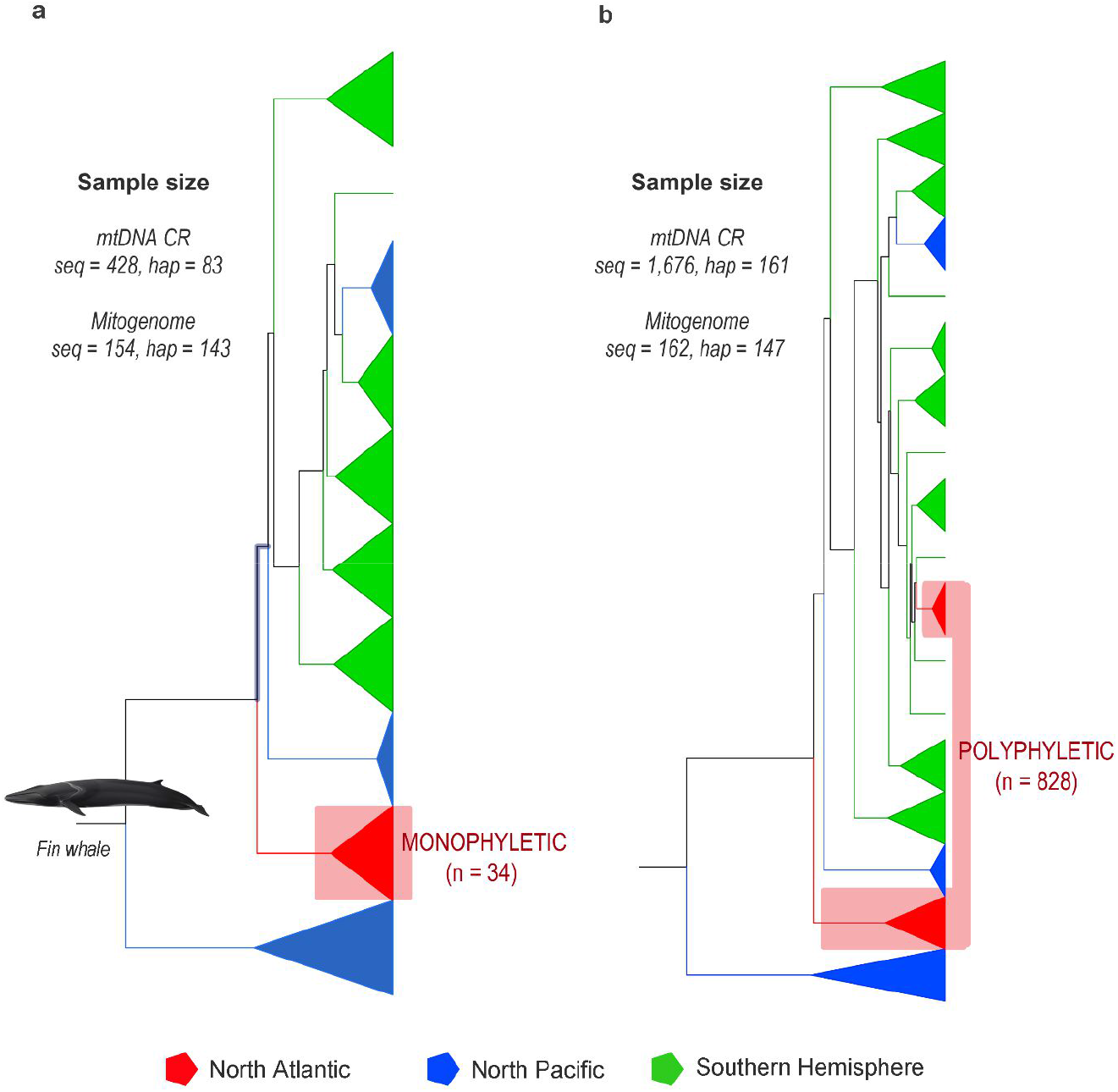

## Introduction

Genealogies estimated from mitochondrial DNA (mtDNA) sequences have been employed towards resolving inter- and intraspecific taxonomic relationships for more than three decades (Avise, 1989; Ball & Avise, 1992; Burbrink *et al*., 2000; Tautz *et al*., 2003; Pons *et al*., 2006). Taxonomic assessments aimed below the nominal species level usually focus on the spatial distinctiveness of monophyletic clades in genealogies estimated from mtDNA sequences, i.e., the presence of phylogeographic structure (Avise *et al*., 1979; Avise *et al*., 1987; Ball & Avise, 1992). The presence of spatially confined monophyletic mitochondrial clades has typically been inferred as evidence for reproductive isolation and consequently some degree of evolutionary distinctiveness. Evolutionary significant units (ESUs) serve as an illustrative example (Ryder, 1986; Bernatchez, 1995). ESUs are generally viewed as distinct components of intraspecific genetic diversity (Ryder, 1986; Bernatchez, 1995). Moritz (1994) proposed that ESUs be defined by the presence of reciprocal monophyly in genealogies estimated from mtDNA sequences as well as “significant” divergence in allele frequencies at nuclear loci between specimens from reciprocally monophyletic clades in the mtDNA sequence genealogy. When monophyly in a mtDNA genealogy is employed as the defining criterion, a key question becomes whether such phylogeographic structure always equates to isolation and evolutionary distinctiveness, and consequently, if the *absence* of monophyly implies a recent common ancestry and evolutionary indistinctiveness. Paetkau (1999) pointed to the fact that the effective population size and time since the most recent common ancestor (TMRCA) are positively correlated. This fundamental relationship implies that isolated populations with a low effective population sizes will become monophyletic at a faster rate compared to populations with larger effective population sizes. This difference has immediate ramifications in those cases when mtDNA monophyly is employed as the main, or sole, criterion in defining ESUs (e.g., Banguera-Hinestroza *et al*., 2002; Lorenzen *et al*., 2008; Archer *et al*., 2013).

Sampling is equally important. Apparent monophyly could be simply a product of insufficient sampling, i.e., an insufficient number specimens to capture all mtDNA clades (Funk & Omland, 2003). Intraspecific genealogies inferred from mtDNA sequences often contain multiple well-supported clades. However, the relative proportions of such clades typically vary across space. Consequently, insufficient sampling in all, or some regions, may result in failure to sample DNA sequences belonging to uncommon clades, erroneously leading to the conclusion of monophyly (Funk & Omland, 2003).

Initially most phylogeographic studies were based solely upon genealogies inferred from mtDNA sequence variation (Avise, 1989; Ball & Avise, 1992; Burbrink *et al*., 2000; Tautz *et al*., 2003; Pons *et al*., 2006). The mtDNA genome (mitogenome) was viewed as especially suitable for this kind of assessments due to its haploid, often maternal and clonal inheritance, which alleviates potential issues in estimating the underlying genealogy from nuclear recombining loci. However, several studies have demonstrated that inferring intraspecific isolation from mtDNA sequences only, could be misleading, ironically because of the maternal inheritance, which prevented detection of male mediated gene flow (Prager *et al*., 1993; Palumbi & Baker, 1994). Consequently, many studies have since complemented mtDNA sequences with nuclear, biparentally-inherited DNA sequences in phylogeographic analyses aimed at detecting evolutionary distinctiveness, such as ESUs as proposed by Moritz (1994).

The relatively recent development of massive parallel sequencing technologies (Funk *et al*., 2012) has led to a resurgence in phylogeographic studies based solely on mtDNA sequences, albeit of the complete mitogenome as opposed to a few hundreds of base pairs (Morin *et al*., 2004; 2010; Archer *et al*., 2013; Meng *et al*., 2013). A search in Web of Science™ (Clarivate Analytics Inc.) revealed that only 14 out of 100 publications aimed at phylogeographic structure or intra-specific taxonomic revisions complemented complete mitogenome sequence data with data from nuclear loci (see Supplementary Materials). The sample sizes in studies based on complete mitogenome sequences in non-model species (Morin *et al*., 2004; Morin *et al*., 2010; Archer *et al*., 2013; Meng *et al*., 2013) remains considerably lower compared to contemporaneous studies based upon Sanger (1981) DNA sequencing of smaller mtDNA regions and nuclear loci (Pastene *et al*., 2007; Halbert *et al*., 2013; Jackson *et al*., 2014). These two aspects, relying solely on mitogenome sequence data (Zachos *et al*., 2013) and low sample sizes, implies that the detection of monophyly is prone to the caveats that haunted earlier, similar studies based upon shorter mtDNA sequences, such as the mtDNA control region (CR). Studies based upon complete mitogenome sequences typically yield very high support for the basal nodes, leading to the impression of high accuracy. However, high accuracy in a single locus genealogy does not necessarily imply that the genealogy accurately reflects the population/subspecies history as has been pointed out by numerous authors in the past (Pamilo & Nei, 1988; Maddison, 1997; Page & Charleston, 1997; Leaché, 2009).

A case in point is *Cetacea* (whales, dolphins and porpoises), a group of highly derived mammals, which has recently been subjected to several re-assessment of species/subspecies status based upon the estimation of intraspecific genealogies from complete mitogenome sequences (Morin *et al*., 2010; Vilstrup *et al*., 2011; Archer *et al*., 2013). The large body sizes, wide ranges and limited availability of osteological specimens in most cetacean species has made it difficult to apply traditional, non-molecular approaches to define intra-specific taxonomic entities and explains the popularity of molecular-based taxonomic assessments in cetaceans. Most baleen whale (*Mysticeti*) species have global distributions and migrate seasonally between low latitude winter breeding grounds and high latitude summer feeding grounds (Ingebrigtsen, 1929; Dawbin, 1966; Jonsgård, 1966; Katona & Whitehead, 1981). As a result, most baleen whale populations roam across entire ocean basins making it challenging to delineate intra-specific evolutionary units. Two aspects are generally assumed, *a priori*, to confine baleen whale distributions and restrict gene flow. The anti-tropical distribution of most baleen whale species presumably acts as a reproductive barrier between the two hemispheres, despite the (proximate) low latitude locations of winter breeding grounds, because the breeding season for each hemisphere is separated by half a year (Davis *et al*., 1998). In addition, most ocean basins are intersected by the continents, which prevent inter-oceanic dispersal as well. Consequently, it is generally assumed that gene flow between con-specific baleen whale populations in different ocean basins is very limited (Valsecchi *et al*., 1997; Bérubé *et al*., 1998; Pastene *et al*., 2007; Morin *et al*., 2010; Jackson *et al*., 2014). Accordingly, current, recognized baleen whale species and subspecies designations typically correspond to ocean basins or hemispheres. For instance, the right whales are comprised of *Eubalaena glacialis*, in the North Atlantic; *E. australis*, in the Southern Hemisphere; and *E. japonica*, in the North Pacific (Rice, 1998; Rosenbaum *et al*., 2000). Similarly Northern Hemisphere blue whales, *Balaenoptera musculus*, are classified as *B. m. musculus* and Southern Hemisphere blue whales as *B. m. intermedia*, in addition to the pygmy blue whale, *B. m. brevicauda* (Rice, 1998).

The fin whale, *Balaenoptera physalus* spp. (Linnaeus, 1758), is a common and globally distributed baleen whale (Gambell, 1985). Fin whales in the Northern Hemisphere are classified as belonging to the subspecies *B. p. physalus* and fin whales in the Southern Hemisphere to *B. p. quoyi* (Fischer, 1829). The fin whale subspecies designations were based upon differences in the vertebrate characteristics (Lönnberg, 1931) as well as traits correlated with body size (Tomilin 1946 cited by Rice, 1998). Employing this classification, North Pacific and North Atlantic fin whales both belong to the same subspecies, despite the observation that gene flow between the two ocean basins is unlikely, at least since the rise of the Panama Isthmus approximately 3.5 million years ago (Coates *et al*., 1992). Recently, Archer and colleagues (2013) employed complete mitogenome sequences from North Atlantic, North Pacific and Southern Hemisphere fin whale specimens to assess the current subspecies status of Northern Hemisphere fin whales. Archer *et al*. (2013) concluded that North Atlantic and some North Pacific fin whales constituted separate subspecies. This conclusion was based upon the observation of a single monophyletic clade that contained all North Atlantic specimens (a sample of 14 specimens), and the presence of several monophyletic clades containing solely North Pacific specimens (Figure 2a). The results of Archer *et al.’s* (2013) mitogenomic analysis appeared to be at odds with previous phylogeographic assessments by Bérubé *et al*. (1998; 2002). Bérubé and co workers based their assessments upon DNA sequences from the highly variable mtDNA CR. Their study identified two mtDNA CR haplotypes in North Atlantic specimens that clustered together with mtDNA CR haplotypes identified among North Pacific specimens. Bérubé and co-workers inferred this result as evidence for recent gene flow between the North Atlantic and North Pacific (Bérubé *et al*., 1998; 2002) likely in a stepping stone manner via the Southern Ocean. In order to resolve the discrepancy between the above-mentioned studies and the support for the proposed taxonomical revision by Archer and colleagues (2013), this study extended the sample size of North Atlantic Ocean (including the Mediterranean Sea) fin whales from the 34 mtDNA CR sequences analyzed by Archer *et al*. (2013) to a total of 786 mtDNA CR sequences. The complete mitogenome was sequenced in a subset (n = 6) of North Atlantic specimens with mtDNA CR haplotypes that clustered with mtDNA CR haplotypes detected in specimens sampled outside the North Atlantic (n = 514). In addition, 20 microsatellite loci were genotyped in 358 specimens from the North Atlantic, North Pacific and Southern Hemisphere. The re-estimation of the genealogy based upon the complete mitogenome sequences from this study showed all ocean basins to be polyphyletic. In other words, these results did not support the current nor the proposed division into subspecies if monophyly in a genealogy estimated from mtDNA sequences is employed as the sole or main defining criterion. The basal topology of the genealogies estimated from the mitogenome and mtDNA CR sequences were qualitatively similar as expected given that the mitogenome represents one linked locus. The assignment test based on the genotype of 20 microsatellite loci revealed that the North Atlantic specimens from the two different clades all belong to the same North Atlantic gene pool. The findings of this study highlight the implications of insufficient sampling when attempting to identify monophyletic clades from mtDNA sequences. However, the results did not negate the possibility that fin whales from different ocean basins could potentially represent different subspecies, although the analysis from this study revealed recent gene flow between fin whales from different ocean basins and hemispheres.

**Figure 1.**
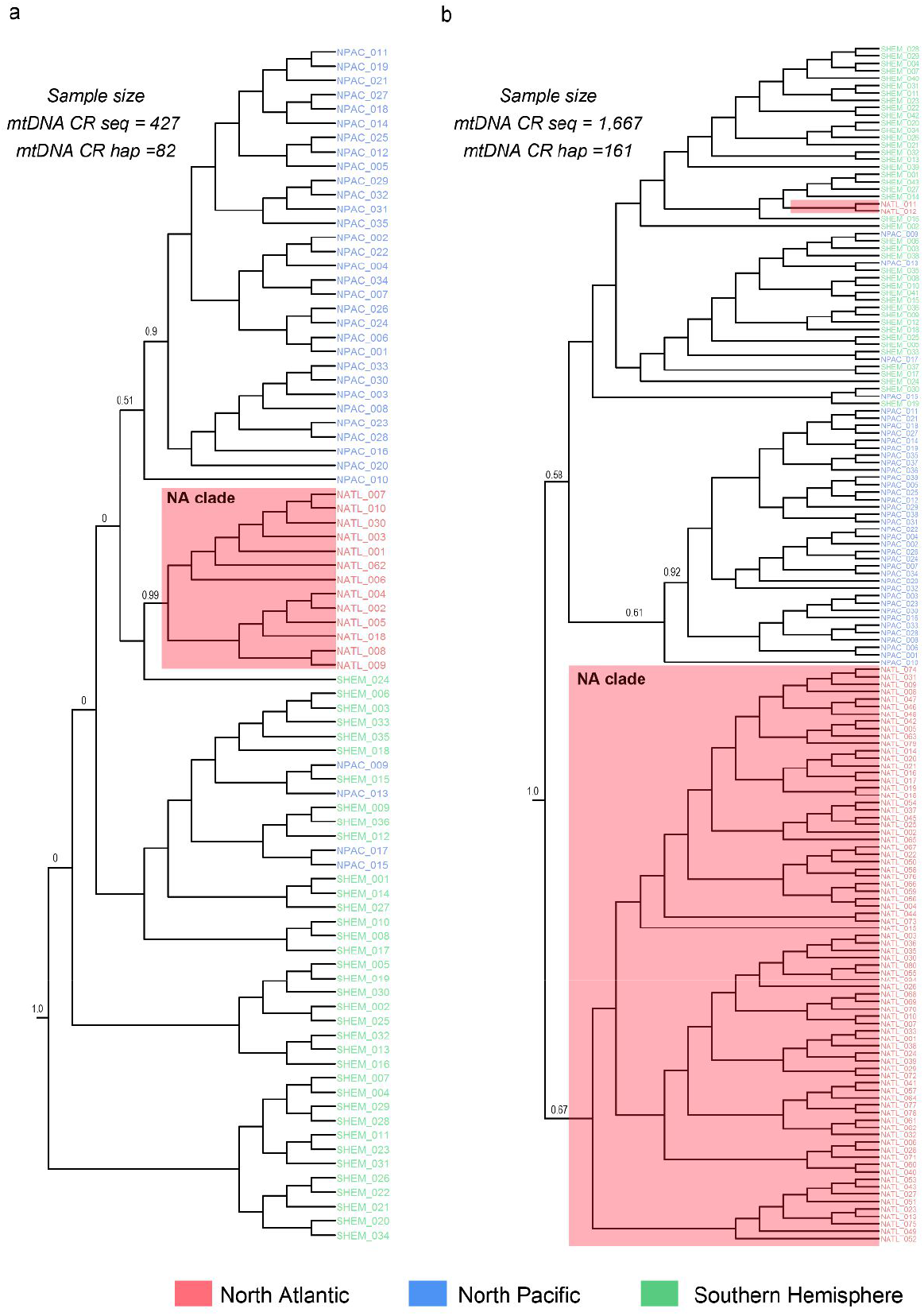
Bayesian genealogy estimated from North Atlantic, North Pacific and Southern Hemisphere fin whale mitochondrial control region (mtDNA CR) haplotypes. Notes: Genealogies were estimated from (a) 82 mtDNA CR haplotypes reported by Archer *et al*. (2013) and (b) 161 mtDNA CR haplotypes reported by Archer *et al*. (2013) combined with additional mtDNA CR haplotypes reported in this study. Colors represent the three ocean basins/regions: the Southern Hemisphere (green, denoted SHEM), the North Pacific (blue, denoted NPAC) and the North Atlantic (red, denoted NATL), respectively. Numbers at basic nodes denotes the posterior probability of the specific node (only the support for basic nodes is reported). A humpback whale (*Megaptera novaeangliae*) mtDNA CR haplotype (Genbank NC_006927) was employed to root the tree (not shown).

**Figure 2.**
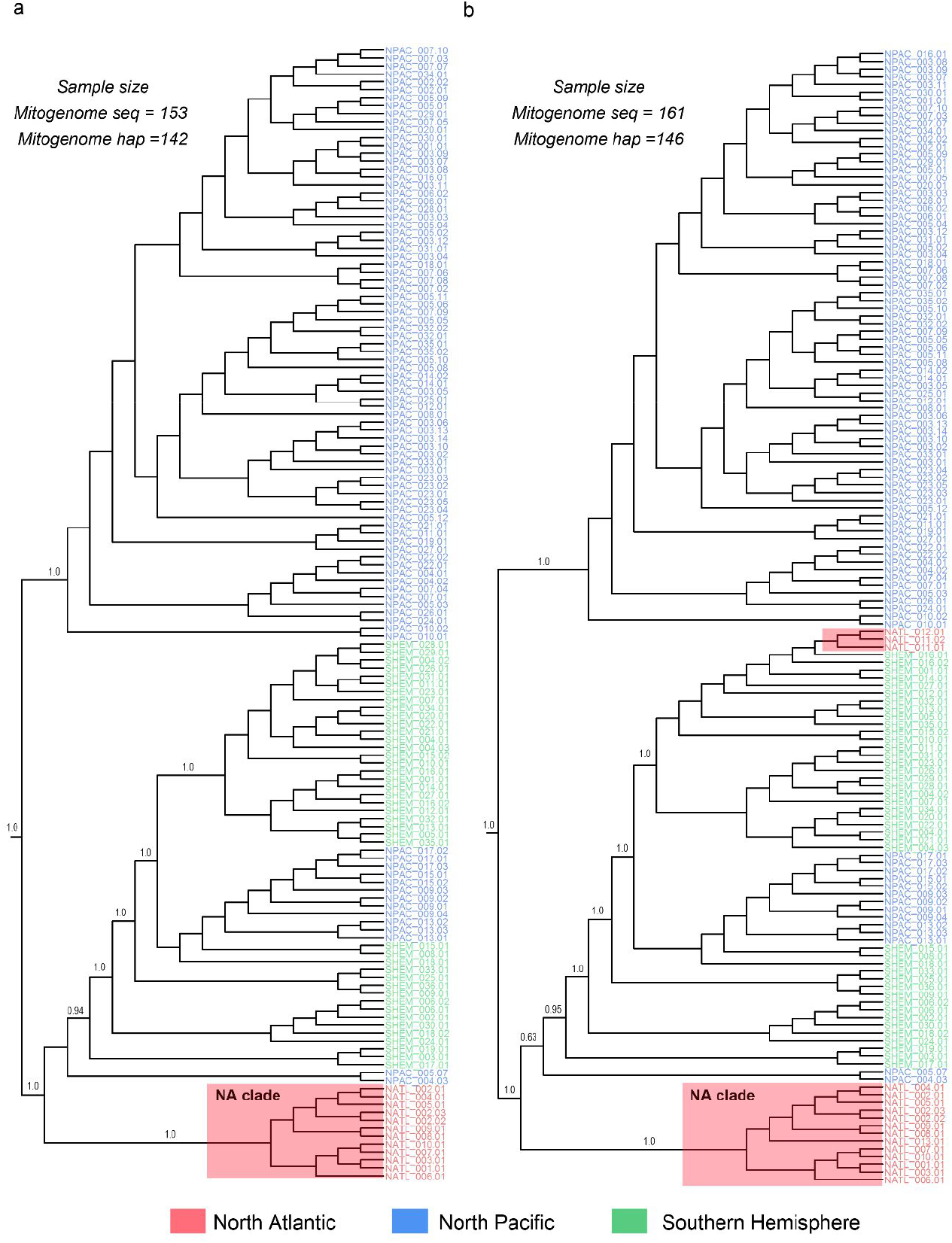
Bayesian genealogy estimated from North Atlantic, North Pacific and Southern Hemisphere fin whale mitochondrial genome (mitogenome) haplotypes. Notes: Genealogies were estimated from (a) 142 mitogenome haplotypes reported by Archer *et al*. (2013) and (b) 146 mitogenome haplotypes reported by Archer *et al*. (2013) combined with additional mitogenome haplotypes reported in this study. Colors represent the three ocean basins/regions: the Southern Hemisphere (green, denoted SHEM), the North Pacific (blue, denoted NPAC) and the North Atlantic Ocean (red, denoted NATL), respectively. Numbers at basic nodes denotes the posterior probability of the specific node (only the support for basic nodes is reported). A humpback whale (*Megaptera novaeangliae*) mitogenome haplotype (Genbank NC_006927) was employed to root the tree (not shown).

More generally, employing monophyly in genealogies based upon DNA sequences from nonrecombining genomes to classify subspecies ignores fundamental population genetic processes as well as key practical issues. These caveats make the approach less valid than its current widespread use suggests. Although these caveats have been highlighted earlier (Paetkau, 1999; Funk & Omland, 2003), the approach has nevertheless regained momentum given the ease of applying massive parallel sequencing technologies to uniparentally inherited, non-recombining genomes, such as the mitogenome.

## Materials and methods

### Sample collection

Tissue samples were obtained from fin whales in the North Atlantic Ocean basin and the Mediterranean Sea (henceforth referred to collectively as the North Atlantic); the North Pacific Ocean basin and the Sea of Cortez (henceforth referred to collectively as the North Pacific) as well as the Southern Hemisphere between 1982 and 2014. Most tissue samples were collected as skin biopsies from free-ranging fin whales as described by Palsbøll *et al*. (1991). The tissue samples originating from Iceland and Spain were collected from whaling operations prior to the international moratorium on commercial whaling. Some samples collected in Greenland originated from local subsistence whaling and some samples collected in US waters originated from dead, beached individuals. All samples were collected in agreement with national and international regulations. Samples were preserved in 5M NaCl with 20% dimethyl sulfoxide and stored at −20 degrees Celsius (Amos & Hoelzel, 1991).

### Mitochondrial DNA sequence data

The mtDNA sequence data were either generated during this study or from data previously published by Archer *et al*. (2013) deposited in the Dryad data repository (http://dx.doi.org/10.5061/dryad.084g8). The experimental methods used to generate the published data were described by Archer *et al*. (2013).

The DNA sequence data generated for this study were obtained in the following manner. Total-cell DNA was extracted from tissue samples either by phenol/chloroform extraction as described by Sambrook and Russell (2001) or using the Qiagen DNAEasy™ Blood and Tissue Kit columns (QIAGEN Inc., Valencia, CA, USA) following the manufacturer’s instructions. Samples were sexed using the ZFY/ZFX multiplexing system as described by Bérubé and Palsbøll (1996b); Bérubé and Palsbøll (1996a). MtDNA CR DNA sequencing was performed as previously described either by (i) Palsbøll *et al*. (1995), but replacing the original reverse primer with BP16071, (Drouot *et al*., 2004); or by (ii) Bérubé *et al*. (2002). The complete mitogenome was sequenced from eight selected specimens. These specimens were selected from genealogy estimated from all mtDNA CR haplotypes. Six specimens were selected among the 33 North Atlantic specimens with mtDNA CR haplotypes that clustered with specimens sampled in other ocean basins. The remaining two specimens were selected among specimens from the North Atlantic and North Pacific Ocean, respectively, with mtDNA CR haplotypes that clustered within monophyletic clades with other specimens sampled in the same ocean. A total of 35 nested primer pairs were employed (Supplementary Materials, Table S1) to amplify and determine the DNA sequence of the complete fin whale mitogenome from partially overlapping ~500 base pair (bp) fragments. PCR (Mullis & Faloona, 1987) amplifications and DNA sequencing were performed under conditions identical to those described above for the mtDNA CR sequencing albeit at different annealing temperatures (Supplementary Materials, Table S1).

### Microsatellite genotyping

The genotype was determined at 20 diploid, autosomal microsatellite loci in samples from four different regions; the North Atlantic (including the Mediterranean Sea) (n=266), the Eastern North Pacific (n=25), the Sea of Cortez (n=46) and the Southern Hemisphere (i.e., the Antarctic) (n=21). The specific microsatellite loci were: AC087, CA234 (Bérubé *et al*., 2005), EV00, EV037, EV094 (Valsecchi & Amos, 1996), GATA028, GATA098, GATA417 (Palsbøll *et al*., 1997), GATA25072, GATA43950, GATA5947654, GATA6063318, GATA91083 (Bérubé et al., in prep), GT011 (Bérubé *et al*., 1998) GT023, GT211, GT271, GT310, GT575, (Bérubé *et al*., 2000) and TAA023 (Palsbøll *et al*., 1997) with tetra, tri-or dimer repeat motifs (Supplementary Materials, Table S2). Individual PCR amplifications were performed in 10 μL volumes, each containing ~ 2-10 ng of extracted DNA, 0.2 μM of each oligonucleotide primer (Supplementary Materials, Table S2) and 1X final QIAGEN Microsatellite PCR Multiplex Mix™ (Qiagen Inc.). Thermo-cycling was carried out on a MJ Research PTC-100™ Thermal Cycler (BioRad Inc.). The PCR amplification consisted of an initial step of five minutes at 95 degrees Celsius, followed by 35 cycles; each of 30 seconds at 95 degrees Celsius, 90 seconds at 57 degrees Celsius and 30 seconds at 72 degrees Celsius. The final step was 10 minutes at 68 degrees Celsius. PCR reactions were diluted 60 times with MilliQ water and then 1μL of diluted PCR reaction was added to 9μL of GeneScan-500™ ROX (Applied Biosystems Inc.) and deionized formamide (GeneScan-500™ ROX 1μL: 70μL) prior to electrophoresis on an ABI 3730™ capillary sequencer (Applied Biosystems Inc.). The length of each amplification product was determined using GeneMapper™ ver. 4.0 (Applied Biosystems Inc.).

### Assembly and analysis of the mitochondrial DNA sequences

MtDNA sequences were aligned and assembled against the fin whale mitogenome sequence deposited in GenBank™ (accession # NC001321) by Árnason *et al*. (1991) using SEQMAN™ (ver. 5.05, DNASTAR Inc.) using default parameter settings. All DNA sequences were trimmed to equal length, i.e., 16,423 and 285 bp for the mitogenome and mtDNA CR DNA sequences, respectively.

#### Estimation of mtDNA haplotype genealogies and divergence times

The genealogies of the mtDNA CR and complete mitogenome haplotypes as well as divergence times were estimated employing the software BEAST (ver. 1.8.2, Drummond & Rambaut, 2007; Drummond *et al*., 2012) largely following the approached by Archer *et al*. (2013). However, in contrast to Archer *et al*. (2013), only a single copy of each haplotype from each ocean basin (both complete and CR mtDNA sequences) was included in each data set. Insertion and deletions were coded as a fifth character. Genealogies were rooted with the homolog DNA sequence from humpback whale, *Megaptera novaeangliae*, (GenBank™ accession #NC006927, (Sasaki *et al*., 2005)) using the alignment reported by Archer *et al*. (2013). The most probable nucleotide mutation model and associated parameter values (e.g., the transition:transversion ratio, the proportion of invariable sites and the gamma distribution) was determined using software JMODELTEST (ver. 2, Darriba *et al*., 2012) and selected using the Bayesian information criterion. The HKY + I + G substitution model (Hasegawa *et al*., 1985) with four substitution categories was selected. A strict molecular clock with a uniform prior distribution and rates between 1 × 10^−5^ and 1 × 10^−2^ substitutions per site per million years was assumed. Similar to Archer *et al*. (2013), the Yule speciation model was employed and the tree topology and branch lengths were initialized with the unweighted pair group method and an arithmetic mean. The TMRCA between the fin whale and humpback whale (Sasaki *et al*., 2005) was employed as a prior for the root of the genealogy, i.e., 15.8-2.8 million years. The prior of the TMRCA was normally distributed. The genealogy and the posterior distribution of the divergence time parameters was estimated using Monte Carlo Markov chains (MCMC) sampling. Samples were drawn every 1,000 steps from a total of 2 × 10^7^ steps of which the first 10 % was discarded as burn-in. Convergence to stationarity and mixing were evaluated with TRACER (ver. 1.5, Rambaut & Drummond, 2007). The consensus genealogy as well as the posterior probability for major nodes and divergence times were obtained using TREEANNOTATOR (ver. 1.8.3), as implemented in BEAST.

#### Estimation of genetic diversity and immigration rates

The software MIGRATE-N (ver. 3.6.6, Beerli & Felsenstein, 1999, 2001) was employed to estimate the genetic diversity (Θ) and immigration rate scaled by the generational mutation rate (*M*) per nucleotide site among the North Atlantic, North Pacific and Southern Hemisphere. The prior ranges of Θ and *M* were determined from preliminary estimations with reduced sample sizes and short MCMC chains with the *F*_st_ -based method as starting values. The prior ranges were subsequently adjusted according to the outcomes of these preliminary estimations, i.e., Θ (uniform prior, min: 0, max: 0.25, *∂*: 0.025) and *M* (uniform prior, min: 0, max: 250.0, *∂*: 25.0). Data sets larger than 100 DNA sequences were subjected to random sub-sampling (without replacement) at sample sizes of 100 DNA sequences per sample partition. Due to significant levels of intra-ocean population structure in mtDNA DNA sequence variation (Bérubé *et al*., 1998; Palsbøll *et al*., 2004; Rivera-León *et al*., under review), the samples from both the Mediterranean Sea and the Sea of Cortez were excluded from the MIGRATE-N estimation among the North Atlantic, North Pacific and Southern Hemisphere. The final estimates were inferred from three independent estimations. Each estimation was initiated with a different random seed and comprised 100 replicates, each consisting of a single long MCMC with 10 million steps discarded as burn in followed by an additional 10 million steps, sampled at every 200^th^ step. A static heating scheme of four chains at temperatures 1.0, 1.5, 3.0, and 1,000,000, respectively, was employed. Convergence was assessed employing the R-CRAN package CODA (ver. 0.19-1, Plummer *et al*., 2006). Consistency among the three independent estimations, smooth and unimodal distribution within the prior range and an effective sample size above 100,000 for all parameter estimates were also considered as indications of convergence (Supplementary Materials, Table S3).

### Multi-locus genotype assignments

The likelihood of multi-locus microsatellite genotypes given the observed allele frequencies in each putative source population was estimated using the probability of identity (Paetkau & Strobeck, 1994) as implemented in GENECLASS v. 2.0 (Piry *et al*., 2004). The null-distribution was estimated from 10,000 multi-locus genotypes drawn at random with replacement from the observed data. The type 1 error rate was set to the default value at 0.01. The observed allele frequencies in each putative source population was estimated from 244 multi-locus microsatellite genotypes for the North Atlantic (including the Mediterranean Sea), 24 for the Eastern North Pacific, 46 for the Sea of Cortez and 21 for the Southern Hemisphere (i.e., the Antarctic).

## Results

The final data sets comprised 1,676 fin whale mtDNA CR DNA sequences and 162 fin whale complete mitogenome sequences (Table 1). Among the final 1,676 mtDNA CR DNA sequences, 428 DNA sequences were obtained from Archer *et al*. (2013) and 1,248 were generated for this study. A total of 410 mtDNA CR sequences from the 782 North Pacific were collected in the Sea of Cortez, a population with low mtDNA sequence diversity (Bérubé *et al*., 2002). Among the 828 North Atlantic mtDNA CR sequences, 115 were collected in the Mediterranean Sea. A total of 161 haplotypes were detected among the 1,676 fin whale mtDNA CR sequences, (Table 1) and 147 haplotypes among the 162 complete mitogenome sequences of which 154 were published by Archer *et al*. (2013) and eight generated during this study (Table 1).

**Table 1.**
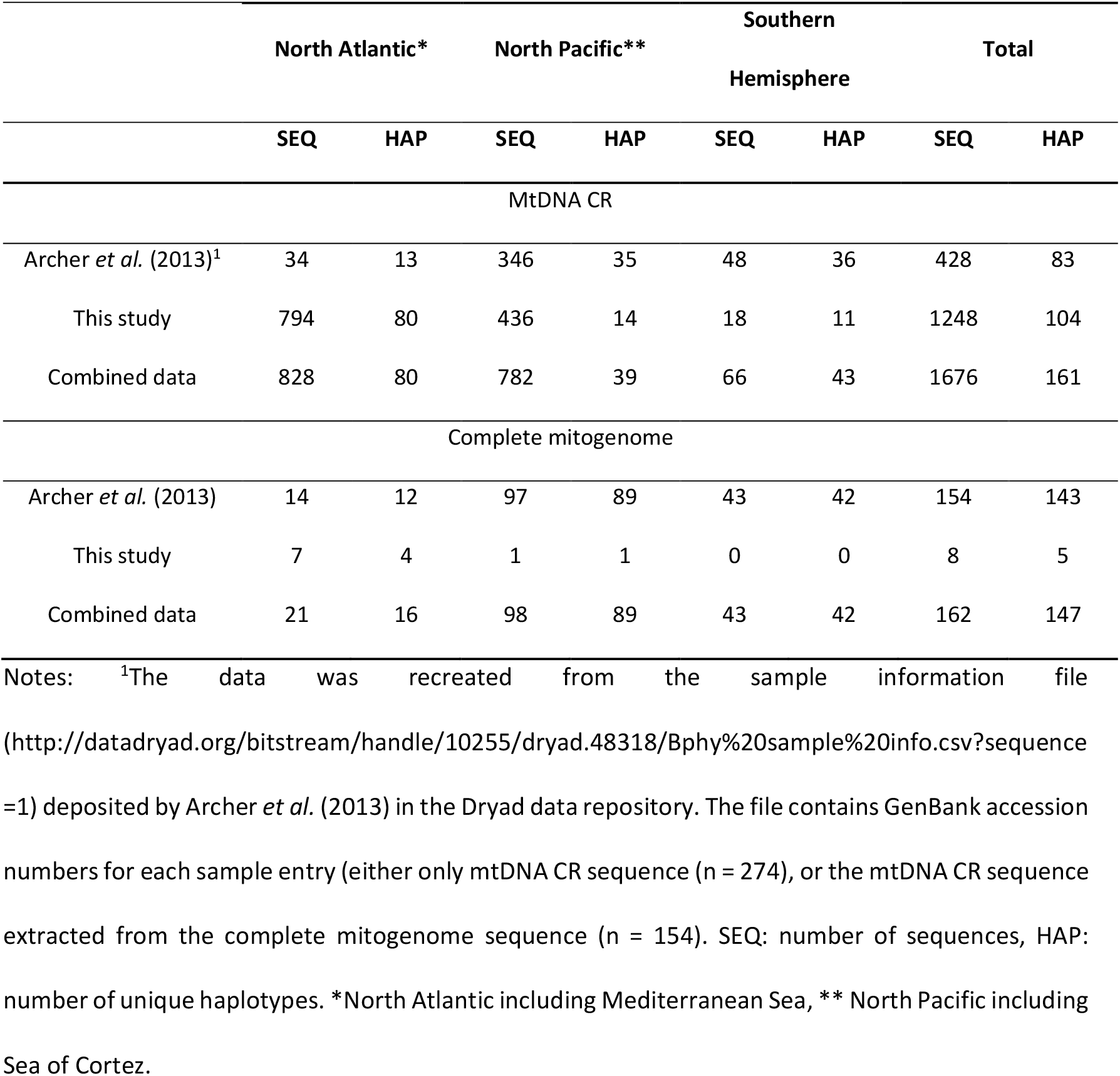
Mitochondrial DNA control region (mtDNA CR) and complete mitochondrial DNA genome (mitogenome) sequences and haplotypes per ocean basin

|  | North Atlantic* |  | North Pacific** |  | Southern Hemisphere |  | Total |  |
| --- | --- | --- | --- | --- | --- | --- | --- | --- |
|  | SEQ | HAP | SEQ | HAP | SEQ | HAP | SEQ | HAP |
| MtDNA CR |  |  |  |  |  |  |  |  |
| Archer *et al.* (2013)<sup>1</sup> | 34 | 13 | 346 | 35 | 48 | 36 | 428 | 83 |
| This study | 794 | 80 | 436 | 14 | 18 | 11 | 1248 | 104 |
| Combined data | 828 | 80 | 782 | 39 | 66 | 43 | 1676 | 161 |
| Complete mitogenome |  |  |  |  |  |  |  |  |
| Archer *et al.* (2013) | 14 | 12 | 97 | 89 | 43 | 42 | 154 | 143 |
| This study | 7 | 4 | 1 | 1 | 0 | 0 | 8 | 5 |
| Combined data | 21 | 16 | 98 | 89 | 43 | 42 | 162 | 147 |

Notes: <sup>1</sup>The data was recreated from the sample information file (http://datadryad.org/bitstream/handle/10255/dryad.48318/Bphy%20sample%20info.csv?sequence=1) deposited by Archer *et al.* (2013) in the Dryad data repository. The file contains GenBank accession numbers for each sample entry (either only mtDNA CR sequence (n = 274), or the mtDNA CR sequence extracted from the complete mitogenome sequence (n = 154). SEQ: number of sequences, HAP: number of unique haplotypes. \*North Atlantic including Mediterranean Sea, \*\* North Pacific including Sea of Cortez.

Both genealogies estimated from the complete mitogenome and mtDNA CR haplotypes published by Archer *et al*. (2013) identified a single monophyletic clade containing all, and only, North Atlantic specimens (denoted NA clade in Figure 1a and 2a). In contrast, haplotypes detected in North Pacific and Southern Hemisphere specimens were polyphyletic (Figure 1a and 2a). In contrast, the genealogies estimated from the complete mitogenome and mtDNA CR haplotypes including both the new data generated in this study and those published by Archer *et al*. (2013) partitioned the North Atlantic specimens in two major clades; one clade (denoted NA clade in Figures 1b) comprised only North Atlantic specimens and another clade comprised DNA sequence haplotypes detected in specimens from the North Pacific and Southern Hemisphere, in addition to the North Atlantic (NATL_011 and NATL_012, Figure 1b). The genealogy estimated from the novel and previously published complete mitogenome haplotypes was similar to the genealogy inferred from the mtDNA CR sequences (Figures 2b). In agreement with the genealogy estimated from the mtDNA CR sequences, North Atlantic specimens were partitioned into two different clades; one clade containing solely North Atlantic specimens (NA clade, Figure 2b) and another clade containing specimens from the North Pacific, Southern Hemisphere and North Atlantic (Figure 2b). The latter clade contained three haplotypes (NATL_011.01, NATL_011.02 and NATL_012.01, Figure 2b) represented by all six North Atlantic specimens from which complete mitogenome DNA sequences were generated during this study (Figure 2b).

The time since the most recent common ancestor (TMRCA) estimated from all complete mitogenome haplotypes included this study (Figure 2b) was estimated at 1.9 million years and the 95% HPD (highest probability density) interval from 1.1 to 2.8 million years (Table 2). The divergence time of the three North Atlantic complete mitogenome haplotype, which clustered outside the NA clade (Figure 2b) detected during this study was estimated at 0.095 million years and the 95% HPD interval from 0.04 to 0.17 million years. The TMRCA for all the complete mitogenome haplotypes detected in the North Atlantic was estimated at 0.99 million years and the 95% HPD from 0.54 to 1.4 million years (Table 2). This estimate was 0.45 million years older than that reported by Archer *et al*. (2013). The TMRCA for all the mtDNA CR haplotypes included in this study (Figure 1b) was estimated at 4.3 million years and the 95% HPD interval from 1.97 to 6.8 million years. In the case of the North Atlantic fin whales, the TMRCA was estimated at 4.2 million years and the 95% HPD interval from 1.96 to 6.8 million years.

**Table 2.**
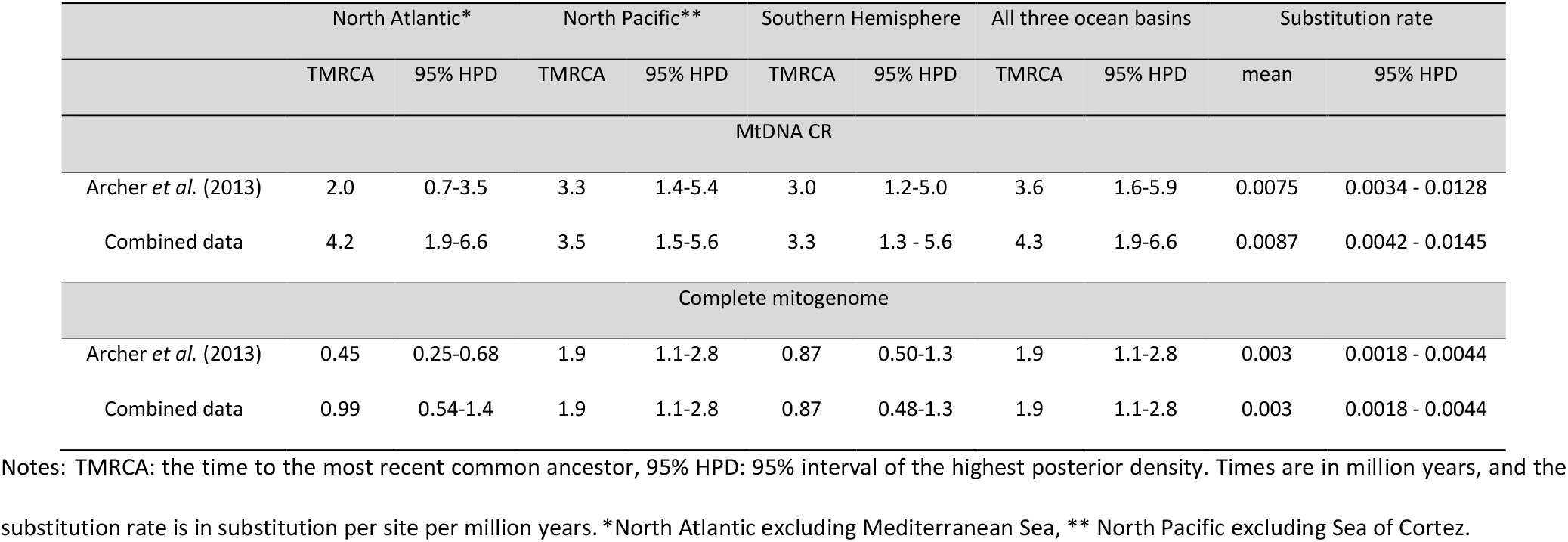
Estimates of time to most recent common ancestor and substitution rates obtained from the mitochondrial DNA control region (mtDNA CR) and complete mitochondrial genome (mitogenome) sequences.

The population origin of 25 fin whale specimens (22 sampled in the North Atlantic, one in the North Pacific, one in the Sea of Cortez and one in the Southern Hemisphere) was inferred from the assignment tests based upon diploid genotypes at 20 microsatellite loci. From the 22 fin whale specimens from the North Atlantic, 20 samples had mtDNA CR haplotypes that were assigned to clades containing specimens from the North Pacific and Southern Hemisphere (i.e., outside the NA clade, Figures 1b). The other two North Atlantic samples were from the main North Atlantic clade (NA clade, Figures 1b). All North Atlantic specimens were assigned to the North Atlantic population. Similarly, the specimen originated from the North Pacific, the specimen originated from the Sea of Cortez and the specimen originated from the Southern Hemisphere (Table 4) were all assigned to their source population, i.e., North Pacific, Sea of Cortez and Antarctic, respectively.

The number of migrants per generation (i.e., *N_e_m* = *ΘM*) estimated from the mtDNA CR sequences from the Southern Hemisphere into the North Pacific was estimated at 0.36 (95% credible interval: 0 - 3.41, Table 3) or one migrant every 2.8 generations. The number of migrants per generation from the Southern Hemisphere into the North Atlantic was similar, i.e., estimated at 0.37 (95% credible interval: 0 - 2.79, Table 3) or one migrant every 2.6 generations. The number of migrants per generation from the North Atlantic into the North Pacific was estimated at 0.0015 (95% credible interval: 0 - 1.56, Table 3) and at 0.0029 (95% credible interval: 0 - 1.97, Table 3) in the opposite direction.

**Table 3.** Average estimates of genetic diversity (*θ*) and number of immigrants per generation (*N_e_m*) for the North Atlantic, North Pacific and Southern Hemisphere^*^

| Parameter | $\theta_{NA}$ | $\theta_{NP}$ | $\theta_{SH}$ | $N_e m_{NP \rightarrow NA}$ | $N_e m_{SH \rightarrow NA}$ | $N_e m_{NA \rightarrow NP}$ | $N_e m_{SH \rightarrow NP}$ | $N_e m_{NA \rightarrow SH}$ | $N_e m_{NP \rightarrow SH}$ |
| --- | --- | --- | --- | --- | --- | --- | --- | --- | --- |
| mode | 0.037 | 0.018 | 0.078 | 0.0029 | 0.3711 | 0.0015 | 0.3565 | 0.0063 | 0.0063 |
| 95% CI | 0.022-0.057 | 0.008-0.032 | 0.048-0.139 | 0-1.973 | 0-2.791 | 0-1.566 | 0-3.406 | 0-3.082 | 0-18.632 |

Notes: NA: North Atlantic, NP: North Pacific, SH: Southern Hemisphere, $\theta$ : genetic diversity, $N_e$ : effective population size, $m$ : immigration rate per generation, → denotes the direction of migration backward in time, 95% CI: 95% credible interval. \*Estimates were based on the mtDNA CR sequences. Samples from Sea of Cortez and the Mediterranean Sea were excluded from the analysis.

**Table 4.** Multi-locus microsatellite genotype probability value (p-value) per putative population and assigned population.

| Sample ID* | CR haplotype number | North Atlantic (n=244) | Antarctica (n=20) | Eastern North Pacific (n=24) | Sea of Cortez (n=45) | # loci | Missing loci | Assigned population |
| --- | --- | --- | --- | --- | --- | --- | --- | --- |
| NAT0009 | NATL_011 | 0.558 | <0.01 | <0.01 | <0.01 | 20 |  | North Atlantic |
| NAT0017 | NATL_011 | 0.400 | 0.045 | <0.01 | <0.01 | 20 |  | North Atlantic |
| NAT0019 | NATL_011 | 0.938 | 0.062 | 0.049 | <0.01 | 20 |  | North Atlantic |
| NAT0024 | NATL_011 | 0.740 | <0.01 | <0.01 | <0.01 | 20 |  | North Atlantic |
| NAT0647 | NATL_011 | 0.938 | 0.062 | 0.049 | <0.01 | 20 |  | North Atlantic |
| NAT0648 | NATL_003 | 0.502 | 0.025 | 0.013 | <0.01 | 19 | GT023 | North Atlantic |
| NAT0001 | NATL_011 | 0.547 | 0.032 | 0.018 | <0.01 | 20 |  | North Atlantic |
| NAT0002 | NATL_011 | 0.692 | 0.011 | <0.01 | <0.01 | 20 |  | North Atlantic |
| NAT0662 | NATL_011 | 0.547 | 0.032 | 0.018 | <0.01 | 20 |  | North Atlantic |
| NAT0003 | NATL_011 | 0.030 | <0.01 | <0.01 | <0.01 | 20 |  | North Atlantic |
| NAT0705 | NATL_011 | 0.394 | <0.01 | <0.01 | <0.01 | 20 |  | North Atlantic |
| NAT0004 | NATL_011 | 0.125 | 0.053 | 0.019 | <0.01 | 20 |  | North Atlantic |
| NAT0706 | NATL_011 | 0.712 | 0.079 | 0.060 | <0.01 | 20 |  | North Atlantic |
| NAT0707 | NATL_016 | 0.064 | <0.01 | <0.01 | <0.01 | 20 |  | North Atlantic |
| NAT0708 | NATL_011 | 0.610 | 0.012 | 0.055 | <0.01 | 20 |  | North Atlantic |
| NAT0005 | NATL_012 | 0.339 | <0.01 | <0.01 | <0.01 | 20 |  | North Atlantic |
| NAT0709 | NATL_011 | 0.223 | <0.01 | <0.01 | <0.01 | 20 |  | North Atlantic |
| NAT0276 | NATL_011 | 0.049 | <0.01 | <0.01 | <0.01 | 20 |  | North Atlantic |
| NAT0710 | NATL_011 | 0.073 | <0.01 | <0.01 | <0.01 | 20 |  | North Atlantic |
| NAT0296 | NATL_011 | 0.213 | <0.01 | <0.01 | <0.01 | 20 |  | North Atlantic |
| NAT0711 | NATL_011 | 0.016 | <0.01 | <0.01 | <0.01 | 20 |  | North Atlantic |
| NAT0712 | NATL_011 | 0.618 | 0.055 | 0.014 | <0.01 | 20 |  | North Atlantic |
| SHE0010 | SHEM_006 | <0.01 | 0.190 | <0.01 | <0.01 | 19 | EV001 | Antarctic |
| NPA0347 | NPAC_009 | <0.01 | 0.068 | 0.938 | <0.01 | 19 | GATA91083 | Eastern North<br>Pacific |
| SOC0172 | NPAC_005 | <0.01 | <0.01 | 0.076 | 0.169 | 20 |  | Sea of Cortez |

Notes: \*The first three letters from the sample ID represent the region where the samples were collected, NAT denotes the North Atlantic Ocean, SHE denotes the Southern Hemisphere, i.e., Antarctic, NPA denotes the Eastern North Pacific Ocean and SOC denotes the Sea of Cortez. The n value in parenthesis represents the sample size for each putative source population. The assigned population was based on the most likely population and its relative score. CR: mtDNA control region.

## Discussion

The initial reason for undertaking this study was the discrepancy between Archer *et al*. (2013) findings and the earlier work published by Bérubé *et al*. (2002; 1998). However, there was a more general concern about the recent resurge in mitogenomic-based studies employing monophyly to delineate intraspecific evolutionary distinct units.

In diploid, recombining genomes, such as the nuclear genome, recombination facilitates that the population-wide variation becomes incorporated into each haploid complement (Pamilo & Nei, 1988). Accordingly, population-specific monophyly at recombining loci requires substantial reproductive isolation for a considerable number of generations. The number of generations depends upon the effective population size and is subject to a large degree of stochasticity (see Hudson & Turelli, 2003). The situation is different for a uniparentally inherited, non-recombining genome, which is sensitive to sampling effects since each lineage contains only the variation of its own lineage rather than the population at large. This fact emphasizes the importance of a sampling scheme that ensures all key clades are sampled if present in the region or putative subspecies. It appears that it is such sampling effect that was the cause for the monophyly of North Atlantic fin whales observed by Archer *et al*. (2013). Archer and colleagues (2013) included a total of 34 North Atlantic fin whale specimens (including Mediterranean Sea specimens) in their analysis, represented by 13 haplotypes (Table 1). In the extended sample of this study, comprising 828 North Atlantic fin whale mtDNA CR sequences represented by 80 haplotypes, 33 sequences (i.e., ~4 %) represented by two haplotypes, were detected with mtDNA CR haplotypes that clustered outside the main North Atlantic clade (NA clade in Figure 1b). The scarcity of North Atlantic fin whales that carry mtDNA haplotypes clustering outside the North Atlantic could be interpreted as the result of a recent dispersal into the North Atlantic and consequently, the fin whales with these rare mtDNA haplotype represent recent immigrants and hence are not part of the North Atlantic gene pool *per se*. However, the analyses of the biparentally inherited microsatellite loci in this study suggested that these individuals were part of the North Atlantic gene pool and unlikely to originate from the North Pacific (both the Sea of Cortez and the eastern North Pacific) nor the Southern Hemisphere. The probability of all these samples’ multi-locus genotypes was higher in the North Atlantic and Mediterranean Sea than in the other ocean basins.

The estimated divergence times among the mtDNA haplotypes (Table 2) were, for some fundamental nodes, considerably older compared to the divergence times reported by Archer *et al*. (2013). The TMRCA among North Atlantic specimens was older due to the polyphyly of the North Atlantic haplotypes in the larger sample. The polyphyly in all three-ocean basins implies that the intra-oceanic TMRCA was similar to the TMRCA for the global sample (Table 2). The issues of low sample and spatially uneven sample sizes in regions outside the North Atlantic have ramifications for any inferences made regarding monophyly in these other regions (Figures 1b and 2b). This general and basic sampling issue makes defining subspecies from mitogenomic data highly problematic since the “distinctiveness” is likely to change with the sampling effort. Consequently, it seems that these “higher level” intra-specific classifications should not be based solely on uniparentally inherited genomes. Such classifications should perhaps be founded upon measures of evolutionary distinctiveness that do not rely upon the “absence” of contradicting observations, e.g., the absence of poly-or paraphyly, which in turn is very sensitive to sampling effort and drift. Possible, and likely more robust, criteria would include the degree of gene flow, time since divergence, or a combination hereof (Hey & Nielsen, 2004; Jackson *et al*., 2014) based upon data from biparentally inherited recombining genomes in conjunction with heritable, non-molecular traits such as ecology and morphology (Crandall *et al*., 2000). However, defining exact quantitative criteria for poorly defined entities, such as, subspecies and ESUs is no simple matter.

Defining species and sub-species is a non-trivial issue and, in many instances, hampering implementation of legal protective measures. In response to such more practical applications, Taylor *et al*. (2017) recently proposed employing population genetic statistics (specifically nucleotide divergence) as a means to delineate sub-species in cetaceans. In principle such objective quantitative genetic criteria, have desirable properties, but most such measures are subject to the same sampling and “rate-of-divergence” issues as monophyly discussed above.

Rare, but occasional, gene flow between baleen whale populations in different hemispheres is possible and appeared to have occurred in humpback whales (Jackson et al., 2014) and the Antarctic minke whale, *Balaenoptera bonaerensis* (Glover et al., 2010). Estimates of migration rate among fin whales from the three ocean basins in this study (Table 3) suggested some gene flow between the Southern Hemisphere and the two Northern Hemisphere ocean basins (i.e., the North Atlantic and the North Pacific). However, the wide 95% credible intervals prevented the exclusion of zero migration. The results were also consistent with those reported by Alter *et al*. (2007), who indicated that the only possible route of gene flow between fin whale populations in different ocean basins in the Northern Hemisphere was through the Southern Hemisphere. Recent historical migration between the Southern Hemisphere and both the North Atlantic and North Pacific, respectively, could explain why some North Pacific and North Atlantic fin whale mtDNA haplotypes clustered within the clade containing most Southern Hemisphere specimens.

The current-accepted classification assigns fin whales from the Northern and Southern Hemisphere to two different subspecies. This taxonomic division implies that North Pacific and North Atlantic fin whales belong to the same subspecies and the Southern Hemisphere fin whales to another subspecies. This taxonomic classification was based upon differences in the vertebrae as well as size differences (Lönnberg, 1931; Tomilin, 1946). The basis of these differences has since been questioned by Perrin *et al*. (2009), who suggested that the different latitudinal origin of the holotypes might explain the size difference (Perrin *et al*., 2009). However, this explanation is difficult to evaluate since the holotype that served as the basis for the differences in the vertebrae described by Lönnberg (1931) was not collected and hence is unavailable. Alternatively, if the Northern Hemisphere populations were founded from the Southern Hemisphere the observed polyphyly could be due to incomplete lineage sorting (Avise *et al*., 1984) as suggested by Pastene *et al*. (2007) in the case of common minke whales, *Balaenoptera acutorostrata*, in the Atlantic Ocean. This appears to be the inference drawn by Archer *et al*. (2013), who emphasizes that the three well supported North Pacific groups (Figure 2a) observed in the genealogy estimated from mitogenome sequences could be the result of incomplete lineage sorting. However, whether the patterns observed in the taxon (i.e., monophyletic, polyphyletic or paraphyletic groups) represent sub-species, as opposed to incomplete lineage sorting, population structure and/or incomplete sampling is still unclear.

Unsurprisingly no qualitative differences between the topologies inferred for the basal part of the mitochondrial genealogies were detected when increasing the data from only 285 bp of mtDNA CR sequence to the complete mitogenome DNA sequences. The general support for individual nodes, especially the most recent nodes, increased with the number of bps per haplotype and hence was substantially higher in genealogy estimated based upon complete mitogenome haplotypes. However, in most cases, the basal nodes are the target of interest in analyses of intraspecific variation aimed at assessing subspecies or ESUs. This observation, together with the obvious need for an increased sampling coverage, suggests that it might be worthwhile to first sequence a limited number of mitogenomes from the extreme parts of the species’ distribution. The mitogenome sequences can then serve as a backbone to identify and subsequently specifically target informative regions, which likely can be sequenced efficiently and at low costs using “standard” Sanger sequencing as proposed by Coulson *et al*. (2006). Such a strategy, as opposed to pyro-sequencing of the entire mitogenome in all specimens, facilitates large sample sizes presumably with minimal loss of phylogenetic signal for the most basic parts of the genealogy.

In conclusion, the present study showed that some of the apparent spatially distinct mtDNA haplotype monophyly reported by Archer and colleagues (2013) was due to a sampling bias. Although untested in this study, the same could well be the case for some of the monophyly detected in other ocean basins. Since monophyly essentially relies upon “absence of evidence” for poly or paraphyly proving monophyly, especially below the species level is difficult and prone to biases. As pointed out be Crandall (2000) identifying sub-species or ESUs solely from genetic data is possibly an over-simplistic perspective and require complementary ecological and morphological data. In principle, genetic data are well-suited to assess divergence times and the degree of reproductive isolation (when gene low is low) but the choice of suitable statistics and appropriate threshold values is no simple task.

## Supporting information

## ACKNOWLEDGEMENTS

We are grateful to Hanne Jørgensen, Anna Sellas, Mary Beth Rew and Christina Færch-Jensen for technical assistance. We thank Drs. P. E. Rosel and K. D. Mullin (U.S. National Marine Fisheries Service Southeast Fisheries Science Center) and members of the U.S. Northeast and Southeast Region Marine Mammal Stranding Network and its response teams, including the International Fund for Animal Welfare, the Marine Mammal Stranding Center, Mystic Aquarium, the Riverhead Foundation for Marine Research and Preservation (K. Durham) and the Marine Mammal Stranding Program of the University of North Carolina Wilmington for access to fin whale samples from the western North Atlantic. We thank Gisli Vikingsson for providing samples. We are indebted to Dr. Eduardo Secchi for facilitating data sharing. Data collection in the Southern Ocean was conducted under research projects *Baleias* (CNPq grants 557064/2009-0 and 408096/2013-6), INTERBIOTA (CNPq 407889/2013-2) and INCT-APA (CNPq 574018/2008-5), of the Brazilian Antarctic Program and a contribution by the research consortium ‘Ecology and Conservation of Marine Megafauna - EcoMega-CNPq’. MAS was supported through a FCT Investigator contract funded by POPH, QREN European Social Fund, and Portuguese Ministry for Science and Education. Data collection in the Azores was funded by TRACE-PTDC/MAR/74071/2006 and MAPCET-M2.1.2/F/012/2011 [FEDER, COMPETE, QREN European Social Fund, and Proconvergencia Açores/EU Program]. Fin whale illustration herein is used with the permission of Frédérique Lucas.

## Contributions to the paper

PJP, MB, JPAH and AAC conceived and designed the study. AA, SGB, SB, DB, AB, HAC, PG, SL, FL, VM, SM, NO, CP, SP, RP, CR, JR, CR, RS, MAS, JU, GV, FWW provided data or sample material, MB, JPAH, CPD, WH, VER-L conducted laboratory analyses. AAC conducted the data analysis with contributions from JPAH and inputs from ES. TO conducted the Web of Science review. AAC, JPAH, PJP and MB drafted the manuscript. All authors read, edited, commented on and approved the final manuscript.

## Data accessibility

All input and output data files for the analysis conducted in this manuscript have been deposited in Datadryad.org under accession: PENDING

## References

Alter, S. E., Rynes, E., & Palumbi, S. R. (2007). DNA evidence for historic population size and past ecosystem impacts of gray whales. Proceedings of the National Academy of Sciences of the United States of America, 104(38), 15162–15167. doi: 10.1073/pnas.0706056104

Amos, B., & Hoelzel, A. R. (1991). Long-term preservation of whale skin for DNA analysis. Report of the International Whaling Commission, Special Issue 13, 99–104.

Archer, F. I., Morin, P. A., Hancock-Hanser, B. L., Robertson, K. M., Leslie, M. S., Bérubé, M., … Taylor, B. L. (2013). Mitogenomic phylogenetics of fin whales (*Balaenoptera physalus spp*.): Genetic evidence for revision of subspecies. PLoS One, 8(5), e63396. doi: 10.1371/journal.pone.0063396

Árnason, U., Gullberg, A., & Widegren, B. (1991). The complete nucleotide sequence of the mitochondrial DNA of the fin whale, *Balaenoptera physalus*. Journal of Molecular Evolution, 33(6), 556–568. doi: 10.1007/bf02102808

Avise, J. C. (1989). Gene trees and organismal histories: A phylogenetic approach to population biology. Evolution, 43(6), 1192–1208. doi: 10.2307/2409356

Avise, J. C., Arnold, J., Ball, R. M., Bermingham, E., Lamb, T., Neigel, J. E., … Saunders, N. C. (1987). Intraspecific phylogeography: The mitochondrial DNA bridge between population genetics and systematics. Annual Review of Ecology and Systematics, 18, 489–522. doi: 10.1146/annurev.ecolsys.18.1.489

Avise, J. C., Giblin-Davidson, C., Laerm, J., Patton, J. C., & Lansman, R. A. (1979). Mitochondrial DNA clones and matriarchal phylogeny within and among geographic populations of the pocket gopher, *Geomys pinetis*. Proceedings of the National Academy of Sciences of the United States of America, 76(12), 6694–6698. doi: 10.1073/pnas.76.12.6694

Avise, J. C., Neigel, J. E., & Arnold, J. (1984). Demographic influences on mitochondrial DNA lineage survivorship in animal populations. Journal of Molecular Evolution, 20(2), 99–105. doi: 10.1007/Bf02257369

Ball, R. M., & Avise, J. C. (1992). Mitochondrial DNA phylogeographic differentiation among avian populations and the evolutionary significance of subspecies. Auk, 109(3), 626–636.

Banguera-Hinestroza, E., Cárdenas, H., Ruiz-García, M., Marmontel, M., Gaitán, E., Vázquez, R., & García-Vallejo, F. (2002). Molecular identification of evolutionarily significant units in the Amazon River Dolphin *Inia sp*. (Cetacea:Iniidae). The Journal of Heredity, 93(5), 312–322.

Beerli, P., & Felsenstein, J. (1999). Maximum-likelihood estimation of migration rates and effective population numbers in two populations using a coalescent approach. Genetics, 152(2), 763–773.

Beerli, P., & Felsenstein, J. (2001). Maximum likelihood estimation of a migration matrix and effective population sizes in n subpopulations by using a coalescent approach. Proceedings of the National Academy of Sciences of the United States of America, 98(8), 4563–4568. doi: 10.1073/pnas.081068098

Bernatchez, L. (1995). A role for molecular systematics in defining evolutionarily significant units in fishes. In J. L. Nielsen (Ed.), Evolution and the Aquatic Ecosystem. Defining Unique Units in Population Conservation (Vol. 17, pp. 114–132). Bethesda, Maryland: American Fisheries Society Symposium.

Bérubé, M., Aguilar, A., Dendanto, D., Larsen, F., Notarbartolo Di Sciara, G., Sears, R., … Palsbøll, P. J. (1998). Population genetic structure of North Atlantic, Mediterranean Sea and Sea of Cortez fin whales, *Balaenoptera physalus* (Linnaeus 1758): analysis of mitochondrial and nuclear loci. Molecular Ecology, 7(5), 585–599. doi: 10.1046/j.1365-294x.1998.00359.x

Bérubé, M., Jørgensen, H., McEwing, R., & Palsbøll, P. J. (2000). Polymorphic di-nucleotide microsatellite loci isolated from the humpback whale, *Megaptera novaeangliae*. Molecular Ecology, 9(12), 2181–2183. doi: 10.1046/j.1365-294X.2000.105315.x

Bérubé, M., & Palsbøll, P. (1996a). Erratum of identification of sex in cetaceans by multiplexing with three ZFX and ZFY specific primers. Molecular Ecology, 5(4), 602–602. doi: 10.1111/j.1365-294X.1996.tb00355.x

Bérubé, M., & Palsbøll, P. (1996b). Identification of sex in cetaceans by multiplexing with three ZFX and ZFY specific primers. Molecular Ecology 5(2), 283–287.

Bérubé, M., Rew, M., Skaug, H., Jørgensen, H., Robbins, J., Best, P., … Palsbøll, P. (2005). Polymorphic microsatellite loci isolated from humpback whale, *Megaptera novaeangliae* and fin whale, *Balaenoptera physalus*. Conservation Genetics, 6(4), 631–636. doi: 10.1007/s10592-005-9017-5

Bérubé, M., Urban, J., Dizon, A. E., Brownell, R. L., & Palsbøll, P. J. (2002). Genetic identification of a small and highly isolated population of fin whales (*Balaenoptera physalus*) in the Sea of Cortez, Mexico. Conservation Genetics, 3(2), 183–190. doi: 10.1023/a:1015224730394

Burbrink, F. T., Lawson, R., & Slowinski, J. B. (2000). Mitochondrial DNA phylogeography of the polytypic North American rat snake (*Elaphe obsoleta*): A critique of the subspecies concept. Evolution, 54(6), 2107–2118. doi: 10.1111/j.0014-3820.2000.tb01253.x

Coates, A. G., Jackson, J. B. C., Collins, L. S., Cronin, T. M., Dowsett, H. J., Bybell, L. M., … Obando, J. A. (1992). Closure of the Isthmus of Panama: The near-shore marine record of Costa Rica and Western Panama. Geological Society of America Bulletin, 104(7), 814–828. doi: 10.1130/00167606

Coulson, M. W., Marshall, H. D., Pepin, P., & Carr, S. M. (2006). Mitochondrial genomics of gadine fishes: implications for taxonomy and biogeographic origins from whole-genome data sets. Genome, 49(9), 1115–1130. doi: 10.1139/G06-083

Crandall, K. A., Bininda-Emonds, O. R. P., Mace, G. M., & Wayne, R. K. (2000). Considering evolutionary processes in conservation biology. Trends in Ecology & Evolution, 15(7), 290–295. doi: 10.1016/S0169-5347(00)01876-0

Darriba, D., Taboada, G. L., Doallo, R., & Posada, D. (2012). jModelTest 2: more models, new heuristics and parallel computing. Nature Methods, 9(8), 772–772.

Davis, W., Fargions, S., May, Leming, T. D., Baumgartner, M. F., Evans, E., … Mullin, K. D. (1998). Physical habitat of cetaceans along the continental slop in the Norrh-Central and Western Gulf of Mexico. Marine Mammal Science, 14(3), 490–507.

Dawbin, W. H. (1966). The seasonal migratory cycle of humpback whales. In K. S. Norris (Ed.), Whales, dolphins and porpoises (pp. 145–170). Berkely, CA: University of California Press.

Drouot, V., Berube, M., Gannier, A., Goold, J. C., Reid, R. J., & Palsboll, P. J. (2004). A note on genetic isolation of Mediterranean sperm whales (*Physeter macrocephalus*) suggested by mitochondrial DNA. Journal of Cetacean Research and Management, 6(1), 29–32.

Drummond, A. J., & Rambaut, A. (2007). BEAST: Bayesian evolutionary analysis by sampling trees. BMC Evolutionary Biology, 7(214). doi: 10.1186/1471-2148-7-214

Drummond, A. J., Suchard, M. A., Xie, D., & Rambaut, A. (2012). Bayesian Phylogenetics with BEAUti and the BEAST 1.7. Molecular Biology and Evolution, 29(8), 1969–1973. doi: 10.1093/molbev/mss075

Funk, D. J., & Omland, K. E. (2003). Species-level paraphyly and polyphyly: Frequency, causes, and consequences, with insights from animal mitochondrial DNA. Annual Review of Ecology Evolution and Systematics, 34, 397–423. doi: 10.1146/annurev.ecolsys.34.011802.132421

Funk, W. C., Mckay, J. K., Hohenlohe, P. A., & Allendorf, F. W. (2012). Harnessing genomics for delineating conservation units. Trends in Ecology & Evolution, 27(9), 489–496. doi: 10.1016/j.tree.2012.05.012

Gambell, R. (1985). Fin whale *Balaenoptera physalus* (Linnaeus, 1758). In S. H. Ridgway & S. R. Harrison (Eds.), Handbook of Marine Mammals (pp. 171–192). London, UK: Academic Press Inc.

Glover, K. A., Kanda, N., Haug, T., Pastene, L. A., Oien, N., Goto, M., … Skaug, H. J. (2010). Migration of Antarctic minke whales to the Arctic. PLoS One, 5(12). doi: 10.1371/journal.pone.0015197

Halbert, K. M. K., Goetze, E., & Carlon, D. B. (2013). High cryptic diversity across the global range of the migratory planktonic copepods *Pleuromamma piseki* and *P. gracilis*. Plos One, 8(10). doi: 10.1371/journal.pone.0077011

Hasegawa, M., Kishino, H., & Yano, T. A. (1985). Dating of the human - ape splitting by a molecular clock of mitochondrial DNA. Journal of Molecular Evolution, 22(2), 160–174. doi: 10.1007/Bf02101694

Hey, J., & Nielsen, R. (2004). Multilocus methods for estimating population sizes, migration rates and divergence time, with applications to the divergence of *Drosophila pseudoobscura* and *D. persimilis*. Genetics, 167(2), 747–760. doi: 10.1534/genetics.103.024182

Hudson, R. R., & Turelli, M. (2003). Stochasticity overrules the “three-times rule”: genetic drift, genetic draft, and coalescence times for nuclear loci versus mitochondrial DNA. Evolution, 57(1), 182–190. doi: 10.1111/j.0014-3820.2003.tb00229.x

Ingebrigtsen, A. (1929). Whales caught in the North Atlantic and other seas. Conseil Permanent International pour l’Exploration de la Mer. Rapports et Proces-Verbaux des Reunions, 56, 123–135.

Jackson, J. A., Steel, D. J., Beerli, P., Congdon, B. C., Olavarría, C., Leslie, M. S., … Baker, C. S. (2014). Global diversity and oceanic divergence of humpback whales (*Megaptera novaeangliae*). Proceedings of the Royal Society of London, Series B: Biological Sciences, 281(1786). doi: 10.1098/rspb.2013.3222

Jonsgård, Å. (1966). The distribution of Balaenopteridae in the north Atlantic Ocean. In K. S. Norris (Ed.), Whales, dolphins and porpoises (pp. 114–124). Berkely, CA: University of California Press

Katona, S. K., & Whitehead, H. P. (1981). Identifying humpaback whales using their natural markings. Polar Record, 20(128), 439–444.

Leaché, A. D. (2009). Species tree discordance traces to phylogeographic clade boundaries in North American fence lizards (Sceloporus). Systematic Biology, 58(6), 547–559. doi: 10.1093/sysbio/syp057

Lönnberg, E. (1931). The Skeleton of Balaenoptera brydei Ö. Olsen: Almqvist & Wiksell.

Lorenzen, E. D., Arctander, P., & Siegismund, H. R. (2008). Three reciprocally monophyletic mtDNA lineages elucidate the taxonomic status of Grant’s gazelles. Conservation Genetics, 9(3), 593–601. doi: 10.1007/s10592-007-9375-2

Maddison, W. P. (1997). Gene trees in species trees. Systematic Biology, 46(3), 523–536. doi: 10.2307/2413694

Meng, X. P., Shen, X., Zhao, N. N., Tian, M., Liang, M., Hao, J., … Zhu, X. L. (2013). Mitogenomics reveals two subspecies in *Coelomactra antiquata* (Mollusca: Bivalvia). Mitochondrial DNA, 24(2), 102–104. doi: 10.3109/19401736.2012.726620

Morin, P. A., Archer, F. I., Foote, A. D., Vilstrup, J., Allen, E. E., Wade, P., … Harkins, T. (2010). Complete mitochondrial genome phylogeographic analysis of killer whales (*Orcinus orca*) indicates multiple species. Genome Research, 20(7), 908–916.

Morin, P. A., Luikart, G., Wayne, R. K., & Grp, S. W. (2004). SNPs in ecology, evolution and conservation. Trends in Ecology & Evolution, 19(4), 208–216. doi: 10.1016/j.tree.2004.01.009

Moritz, C. (1994). Defining ‘Evolutionarily Significant Units’ for conservation. Trends in Ecology & Evolution, 9(10), 373–375. doi: 10.1016/0169-5347(94)90057-4

Mullis, K. B., & Faloona, F. A. (1987). Specific synthesis of DNA in vitro via a polymerase-catalyzed chain reaction. Methods in Enzymology, 155, 335–350. doi: 10.1016/0076-6879(87)55023-6

Paetkau, D. (1999). Using genetics to identify intraspecific conservation units: a critique of current methods. Conservation Biology, 13(6), 1507–1509. doi: 10.1046/j.1523-1739.1999.98507.x

Paetkau, D., & Strobeck, C. (1994). Microsatellite analysis of genetic variation in black bear populations. Molecular Ecology, 3(5), 489–495. doi: 10.1111/j.1365-294X.1994.tb00127.x

Page, R. D. M., & Charleston, M. A. (1997). From gene to organismal phylogeny: Reconciled trees and the gene tree species tree problem. Molecular Phylogenetics and Evolution, 7(2), 231–240. doi: 10.1006/mpev.1996.0390

Palsbøll, P. J., Bérubé, M., Aguilar, A., Notarbartolo-Di-Sciara, G., & Nielsen, R. (2004). Discerning between recurrent gene flow and recent divergence under a finite-site mutation model applied to North Atlantic and Mediterranean Sea fin whale (*Balaenoptera physalus*) populations. Evolution, 58(3), 670–675. doi: 10.1111/j.0014-3820.2004.tb01691.x

Palsbøll, P. J., Bérubé, M., Larsen, A. H., & Jørgensen, H. (1997). Primers for the amplification of tri- and tetramer microsatellite loci in baleen whales. Molecular Ecology, 6(9), 893–895. doi: 10.1046/j.1365-294X.1997.d01-214.x

Palsbøll, P. J., Clapham, P. J., Mattila, D. K., Larsen, F., Sears, R., Siegismund, H. R., … P., A. (1995). Distribution of mtDNA haplotypes in North Atlantic humpback whales: the influence of behaviour on population structure. Marine Ecology Progress Series, 116, 1–10.

Palsbøll, P. J., Larsen, F., & Hansen, E. S. (1991). Sampling of skin biopsies from free-ranging large cetaceans in West Greenland: Development of new biopsy tips and bolt designs. Report of the International Whaling Commission (Special Issue 13), 71–79.

Palumbi, S. R., & Baker, C. S. (1994). Contrasting population structure from nuclear intron sequences and mtDNA of humpback whales. Molecular Biology and Evolution, 11(3), 426–435.

Pamilo, P., & Nei, M. (1988). Relationships between gene trees and species trees. Molecular Biology and Evolution, 5(5), 568–583.

Pastene, L. A., Goto, M., Kanda, N., Zerbini, A. N., Kerem, D., Watanabe, K., … Palsbøll, P. J. (2007). Radiation and speciation of pelagic organisms during periods of global warming: the case of the common minke whale, *Balaenoptera acutorostrata*. Molecular Ecology, 16(7), 1481–1495. doi: 10.1111/j.1365-294X.2007.03244.x

Perrin, W. F., Mead, J. G., & Brownell Jr, R. L. (2009). Review of the evidence used in the description of currently recognized cetacean subspecies NOAA Technical Memorandum NMFS - SWFSC- 450.

Piry, S., Alapetite, A., Cornuet, J. M., Paetkau, D., Baudouin, L., & Estoup, A. (2004). GENECLASS2: A software for genetic assignment and first-generation migrant detection. Journal of Heredity, 95(6), 536–539. doi: 10.1093/jhered/esh074

Plummer, M., Best, N., Cowles, K., & Vines, K. (2006). CODA: Convergence diagnosis and output analysis for MCMC. R News, 6, 7–11.

Pons, J., Barraclough, T. G., Gomez-Zurita, J., Cardoso, A., Duran, D. P., Hazell, S., … Vogler, A. P. (2006). Sequence-based species delimitation for the DNA taxonomy of undescribed insects. Systematic Biology, 55(4), 595–609. doi: 10.1080/10635150600852011

Prager, E. M., Sage, R. D., Gyllensten, U., Thomas, W. K., Hubner, R., Jones, C. S., … Wilson, A. C. (1993). Mitochondrial DNA sequence diversity and the colonization of Scandinavia by house mice from East Holsten. Biological Journal of the Linnean Society, 50(2), 85–122.

Rambaut, A., & Drummond, A. J. (2007). Tracer v1. 4.

Rice, C. W. (1998). Marine mammals of the world, systematics and distributuion: Society for Marine Mammalogy Special Publications 4.

Rosenbaum, H. C., Brownell, R. L., Brown, M. W., Schaeff, C., Portway, V., White, B. N., … DeSalle, R. (2000). World-wide genetic differentiation of *Eubalaena:* questioning the number of right whale species. Molecular Ecology, 9(11), 1793–1802. doi: 10.1046/j.1365-294x.2000.01066.x

Ryder, O. A. (1986). Species conservation and systematics: The dilemma of subspecies. Trends in Ecology & Evolution, 1(1), 9–10. doi: 10.1016/0169-5347(86)90059-5

Sambrook, J., & Russell, D. W. (2001). Molecular cloning: a laboratory manual (Third ed.). New York, USA: Cold Spring Harbor Laboratory Press, Cold Spring Harbor.

Sanger, F. (1981). Determination of nucleotide sequences in DNA. Science, 214(4526), 1205–1210. doi: 10.1126/science.7302589

Sasaki, T., Nikaido, M., Hamilton, H., Goto, M., Kato, H., Kanda, N., … Okada, N. (2005). Mitochondrial phylogenetics and evolution of mysticete whales. Systematic Biology, 54(1), 77–90. doi: 10.1080/10635150590905939

Tautz, D., Arctander, P., Minelli, A., Thomas, R. H., & Vogler, A. P. (2003). A plea for DNA taxonomy. Trends in Ecology & Evolution, 18(2), 70–74. doi: 10.1016/S0169-5347(02)00041-1

Taylor, B. L., Archer, F. I., Martien, K. K., Rosel, P. E., Hancock-Hanser, B. L., Lang, A. R., … Baker, C. S. (2017). Guidelines and quantitative standards to improve consistency in cetacean subspecies and species delimitation relying on molecular genetic data. Marine Mammal Science, 33, 132–155. doi: 10.1111/mms.12411

Tomilin, A. G. (1946). Thermoregulation and the geographical races of cetaceans (Termoregulyatsiya I geograficheskie racy kitoobraznykh.). Doklady Akad Nauk CCCP, 54, 465–472.

Valsecchi, E., & Amos, W. (1996). Microsatellite markers for the study of cetacean populations. Molecular Ecology, 5(1), 151–156. doi: 10.1111/j.1365-294X.1996.tb00301.x

Valsecchi, E., Palsbøll, P., Hale, P., GlocknerFerrari, D., Ferrari, M., Clapham, P., … Amos, B. (1997). Microsatellite genetic distances between oceanic populations of the humpback whale (*Megaptera novaeangliae*). [Article]. Molecular Biology and Evolution, 14(4), 355–362.

Vilstrup, J. T., Ho, S. Y. W., Foote, A. D., Morin, P. A., Kreb, D., Krutzen, M., … Gilbert, M. T. P. (2011). Mitogenomic phylogenetic analyses of the Delphinidae with an emphasis on the Globicephalinae. BMC Evolutionary Biology, 11(65). doi: 10.1186/1471-2148-11-65

Zachos, F. E., Apollonio, M., Barmann, E. V., Festa-Bianchet, M., Gohlich, U., Habel, J. C., … Suchentrunk, F. (2013). Species inflation and taxonomic artefacts-A critical comment on recent trends in mammalian classification. Mammalian Biology, 78(1), 1–6. doi: 10.1016/j.mambio.2012.07.083

