## Supplementary material for "Fin whale (*Balaenoptera physalus*) mitogenomics: A cautionary tale of defining sub-species from mitochondrial sequence monophyly"

**Table S1. Oligo-nucleotides employed to amplify and sequence the complete fin whale mitochondrial genome**

| Oligo ID | Nucleotide sequence (5' – 3') | Annealing temperature |
|---|---|---|
| Bp_134F<br>Bp_777R | AGCTGGGCCTGGATGTATTTGT<br>AACACTCTTTACGCCGTGCTTC | 58.2 |
| Bp_396F<br>Bp_1194R | ACATGAACGCCATCCCTATCCA<br>TAGCCCATTACTTCCCAACCCA | 57.55 |
| Bp_781F<br>Bp_1531R | CACGGCGTAAAGAGTGTTAAGG<br>GGGCTAGGGCTAGTTCAAGATA | 55.6 |
| Bp_1388F<br>Bp_2075R | GTTCGTTAACTCAGGCCAAGCA<br>GTTGTCGAGCTTGAACGCTT | 57 |
| Bp_1776F<br>Bp_2500R | TTACCCGAAACCAGACGAGCTA<br>CGTGTGGCCATTCATACAAGTC | 56.95 |
| Bp_2206F<br>Bp_2874R | CTCCTAGCACAAGCTTACACCA<br>CTGCACCATTAGGATGTCCTGA | 56.3 |
| Bp_2684F<br>Bp_3403R | AATTTCGGTTGGGGTGACCT<br>GGGCTAGTACTGGTGCAATGAT | 56.3 |
| Bp_3166F<br>Bp_3858R | GGTTCAAATCCTCTCCCCAACA<br>TTGGCGTATTCTGCCAGGAAG | 56.85 |
| Bp_3575F<br>Bp_4299R | TAATTGGAGCCCTACGAGCAGT<br>GTATGGGCCCGATAGCTTGTTT | 57.3 |
| Bp_3914F<br>Bp_4660R | TCCTAGGAACATTCCACAACCC<br>GGCTAATCCCAGTTTGATGGCT | 56.5 |
| Bp_4508F<br>Bp_5237R | GCCACAGAAGCTTCTACCAAGT<br>GTGCTGTGGAGTAAGTAAGACG | 56.05 |
| Bp_4843F<br>Bp_5604R | AATAGGAGGCTGAGGTGGACT<br>GCGGGAGAAGTAGATTGAAGC | 55.9 |
| Bp_5412F<br>Bp_6191R | ACCCAGACCAAGAGCCTTCAAA<br>TCAACTGAGGCTCCTGCATGTG | 58.65 |
| Bp_5845F<br>Bp_6605R | TTCGGTGCTTGAGCAGGAAT<br>AACCCAATGGACACCATAGCTC | 56.65 |
| Bp_6388F<br>Bp_7105R | GCAGCCGGAATTACAATGCTA<br>TTGTGTAGGCGTCTGGGTAATC | 56.05 |
| Bp_6826F<br>Bp_7501R | ACAGTAGGCGGTCTAACTGGTA<br>TGATGGGTGATGCTGCATCT | 56.3 |
| Bp_7304F<br>Bp_7989R | CAGCATTCGTCAACCCAAAGTG<br>TTAGGCGTCCTGGGATTGCAT | 57.7 |
| Bp_7740F<br>Bp_8479R | TCAATAACCCCTCCCTCACTGT<br>TTAGTCGATTTGGTGCAGGGAA | 56.5 |
| Bp_8100F<br>Bp_8829R | TCCTAGAACTAGTACCCCTAGA<br>CCGGTTGAATGAATAGACTG | 51.25 |

| | | |
|---|---|---|
| Bp_8689F<br>Bp_9398R | AACGTAGGAATGGCTATCCC<br>GTCAACATCCGCCTAGTTCT | 53.7 |
| Bp_9096F<br>Bp_9770R | ATAGTAAACCCCAGCCCTTGAC<br>GTATCAAGCGGCACGTTCAAAG | 56.65 |
| Bp_9571F<br>Bp_10300R | CACTAGGCCTCTACTTCACCTT<br>TCACAGTCTAGTGGGTCGAA | 54.75 |
| Bp_9971F<br>Bp_10699R | CTCGTATTCATCGCCTTCTGAC<br>CTGTGGGCTGTGGAGTTAATTC | 55.55 |
| Bp_10524F<br>Bp_11301R | CCTCCTAGTTTTCGCAGCTTGT<br>CAAGGACTATGGAGCCTGCAAT | 57.05 |
| Bp_10997F<br>Bp_11634R | AGCCACACTAATCCCTACCCTT<br>ATGGTTCGGCTATGAATGCG | 56.15 |
| Bp_11515F<br>Bp_12328R | TCGTCATCGCAGCTATCCTC<br>GACTAGTGATGAAGGCGCAGAA | 56.45 |
| Bp_11986F<br>Bp_12732R | TCATCTTAGGCCCTCTCTACTG<br>GAGTCCAATGTCTCCGATACGA | 55.15 |
| Bp_12526F<br>Bp_13294R | TATATGCACTCCGACCCCTACA<br>TGGTGAATGGGAGGGCCTTAAA | 57.2 |
| Bp_13029F<br>Bp_13798R | AACAGTAACCCTCTGCTTAGGC<br>GGGGTAGGCGATGTATGATTGT | 56.35 |
| Bp_13592F<br>Bp_14290R | TCTTCGCTGGCTTCATCCTA<br>GAACAGTATCCTGAGGTTTGGG | 54.95 |
| Bp_13983F<br>Bp_14746R | CATCACCATCACCCTCAGCATA<br>GCTAGGAATAGGCCTGTTAGGA | 55.65 |
| Bp_14546F<br>Bp_15239R | CACATGGACTTCAACCATGACC<br>TCTATGTCGGATGGGATGCCT | 56.4 |
| Bp_15031F<br>Bp_15854R | TCTGAGGCGCAACTGTAATCAC<br>GTAGAACTTCAGCTTTGGGTGC | 56.8 |
| Bp_15534F<br>Bp_16193R | CACACATCCAATCAACGAAGCA<br>GTGATCTAATGGAGCGGCCATA | 56.3 |
| Bp_16009F<br>Bp_16714R | GCGTCTTTCCATGGGTATGAAC<br>ATCTAGGGACGAGCCTGTCT | 56.7 |

The number after “Bp_” denotes the position of the 3’ end of the oligo in the fin whale genome deposited by Árnason *et al.* (1991) in GenBank™ (accession # NC001321). The suffix, F or R, denotes a forward or reverse oriented oligo, respectively.

**Table S2. Microsatellite loci employed in this study.**

| Locus | Oligo-nucleotide primer sequence (5' - 3') | Range base pairs | Source |
|---|---|---|---|
| GT211 | CAT CTG TGC TTC CAC AAG CCC<br>GGC ACA AGT CAG TAA GGT AGG | 187-213 | Bérubé *et al.* (2000) |
| GT011 | CAT TTT GGG TTG GATCAT TC<br>GTG GAG ACC AGG GAT ATT G | 100-126 | Bérubé *et al.* (1998) |
| GATA25072 | TGG ACA CAT TTA AGG GGA TAA<br>CAA AGT AAG GCG AAT CAA GTT | 377-415 | Bérubé *et al.,* unpublished |
| EV037 | AGC TTG ATT TGG AAG TCA TGA<br>TAG TAG AGC CGT GAT AAA GTG C | 193-223 | Valsecchi and Amos (1996) |
| GATA028 | AAA GAC TGA GAT CTA TAG TTA<br>CGC TGA TAG ATT AGT CTA GG | 290-350 | Palsbøll *et al.* (1997) |
| GATA098 | TGT ACC CTG GAT GGA TAG ATT<br>TCA CCT TAT TTT GTC TGT CTG | 85-129 | Palsbøll *et al.* (1997) |
| GATA417 | CTG AGA TAG CAG TTA CAT GGG<br>TCT GCT CAG GAA ATT TTC AAG | 220-308 | Palsbøll *et al.* (1997) |
| GT575 | CTG CCA ATA TAA GTG AAT ACA<br>CCA TCA ACT GGA AGT CTT T | 137-161 | Bérubé *et al.* (2000) |
| GATA43950 | TGT GGA GAA GAT GGG AAA TAA<br>GTT GTG GGT GAA ATG TTT AGG | 425-449 | Bérubé *et al.,* unpublished |
| TAA023 | CTC GCA CAG AAA TGA AGA CCC<br>AGA GCC TGA ACC AGA ACA AGG | 72-101 | Palsbøll *et al.* (1997) |
| GT271 | GCT CAC ACT GGT AAT CTG TGG<br>CCC TAG GAA GGA TAG ACA TAG | 104-128 | Bérubé *et al.* (2000) |
| EV094 | ATC GTA TTG GTC CTT TTC TGC<br>TGA AGT AAC AGT TAA TAT ACC | 210-238 | Valsecchi and Amos (1996) |
| GATA97408 | AAT GAA CCA ATG GAA CAC AAC<br>CAT GTC GGT CTT TAA TCC ATC | 300-308 | Bérubé *et al.,* unpublished |
| GATA6063318 | GTC CTG AAG AAG GAC TTA GGG<br>GCA CTT AGG CAT CTG GAA GT | 298-326 | Bérubé et al., unpublished |
| AC087 | GCT TCC AGA AGC AAT GAT GGA<br>ACC AGG GTG GGT TCT TAA ACT A | 152-168 | Bérubé *et al.* (2005) |
| GT023 | CAG GAA TCT CAG GGG ATT TA<br>CTT CAG GCA ACA TTT CAT TTT | 106-140 | Bérubé *et al.* (2000) |
| GATA91083 | CCA AAT TGA GAC AGC AAC TCT<br>GTG ATC CTT CTC CTT TCC AAT | 180-220 | Bérubé *et al.,* unpublished |
| CA234 | CAA CCT TAT TCT TGA CCT CAT<br>TGG ATC CTC TAC CTA CCT TAG | 196-220 | Bérubé *et al.* (2005) |
| GT310 | GAA TAC TCC CAG TAG TTT CTC<br>TAA CTT GTG GAA GAT GCC AAC | 100-132 | Bérubé *et al.* (2000) |
| EV001 | GCT GGG GAC ACA AAC AAA AGC TC<br>ATA AAC TCT AAT ACA CTT CCT CCA AC | 168-200 | Valsecchi and Amos (1996) |
| GATA5947654 | CAA AGC ATA AAA CCA GCA ACT<br>GCA TAA GCC AAT TCC TGA TAA | 320-350 | Bérubé *et al.,* unpublished |

**Table S3. Estimates of genetic diversity ($\theta$) and immigration rates ($M$) for the North Atlantic, North Pacific and Southern Hemisphere for the three replicates***

| Replicate 1 | | | | | | | |
|---|---|---|---|---|---|---|---|
| Parameter | 2.50% | 25.00% | mode | 75.00% | 97.50% | median | mean |
| $\theta_{NA}$ | 0.028 | 0.037 | 0.044 | 0.051 | 0.066 | 0.046 | 0.046 |
| $\theta_{NP}$ | 0.009 | 0.015 | 0.020 | 0.024 | 0.034 | 0.021 | 0.021 |
| $\theta_{SH}$ | 0.048 | 0.067 | 0.078 | 0.092 | 0.137 | 0.084 | 0.088 |
| $M_{NP \to NA}$ | 0 | 0 | 0.08 | 7.17 | 29.83 | 7.25 | 9.99 |
| $M_{SH \to NA}$ | 0 | 3.67 | 9.25 | 17.5 | 42.5 | 14.92 | 17.49 |
| $M_{NA \to NP}$ | 0 | 0 | 0.08 | 10.17 | 45.0 | 10.25 | 14.74 |
| $M_{SH \to NP}$ | 0 | 5.33 | 20.25 | 40.83 | 108.17 | 36.25 | 43.41 |
| $M_{NA \to SH}$ | 0 | 0 | 0.08 | 4.83 | 20.5 | 4.92 | 6.66 |
| $M_{NP \to SH}$ | 0 | 0 | 0.08 | 9.0 | 133.83 | 9.08 | 26.37 |
| Replicate 2 | | | | | | | |
| Parameter | 2.50% | 25.00% | mode | 75.00% | 97.50% | median | mean |
| $\theta_{NA}$ | 0.022 | 0.031 | 0.036 | 0.043 | 0.056 | 0.038 | 0.039 |
| $\theta_{NP}$ | 0.0052 | 0.010 | 0.014 | 0.017 | 0.025 | 0.015 | 0.015 |
| $\theta_{SH}$ | 0.048 | 0.067 | 0.079 | 0.093 | 0.149 | 0.085 | 0.091 |
| $M_{NP \to NA}$ | 0 | 0 | 0.08 | 8.17 | 34 | 8.25 | 11.4 |
| $M_{SH \to NA}$ | 0 | 4.00 | 10.08 | 19.83 | 48.5 | 17.08 | 19.92 |
| $M_{NA \to NP}$ | 0 | 0 | 0.08 | 12.67 | 57.33 | 12.75 | 18.63 |
| $M_{SH \to NP}$ | 0 | 5.83 | 19.58 | 41.17 | 113.5 | 36.42 | 44.62 |
| $M_{NA \to SH}$ | 0 | 0 | 0.08 | 5.0 | 21.67 | 5.08 | 7.01 |
| $M_{NP \to SH}$ | 0 | 0 | 0.08 | 8.67 | 147.67 | 8.75 | 29.7 |
| Replicate 3 | | | | | | | |
| Parameter | 2.50% | 25.00% | mode | 75.00% | 97.50% | median | mean |
| $\theta_{NA}$ | 0.018 | 0.025 | 0.030 | 0.035 | 0.048 | 0.032 | 0.032 |
| $\theta_{NP}$ | 0.011 | 0.017 | 0.022 | 0.026 | 0.037 | 0.023 | 0.023 |
| $\theta_{SH}$ | 0.049 | 0.067 | 0.078 | 0.092 | 0.133 | 0.084 | 0.087 |
| $M_{NP \to NA}$ | 0 | 0 | 0.08 | 9.5 | 40.33 | 9.58 | 13.47 |
| $M_{SH \to NA}$ | 0 | 3.83 | 10.92 | 22.0 | 56.33 | 19.08 | 22.67 |
| $M_{NA \to NP}$ | 0 | 0 | 0.08 | 10.0 | 44.5 | 10.08 | 14.48 |
| $M_{SH \to NP}$ | 0 | 4.33 | 18.42 | 36.0 | 97.67 | 32.25 | 38.89 |
| $M_{NA \to SH}$ | 0 | 0 | 0.08 | 5.17 | 24.17 | 5.25 | 7.67 |
| $M_{NP \to SH}$ | 0 | 0 | 0.08 | 8.33 | 119.5 | 8.42 | 23.57 |

Notes: NA: North Atlantic, NP: North Pacific, SH: Southern Hemisphere, $\theta$: genetic diversity, $M$: immigration rate, → denotes the direction of migration forward in time. *Samples from Sea of Cortez and the Mediterranean Sea were excluded from the analysis. Estimates of number of migrants per generation ($N_e m$ ) enlisted on Table 3 were estimated as $\Theta_i M_{j \to i}$.

### Literature survey

**Studies based on mitochondrial genome data complemented with nuclear data**

How many published studies complemented phylogenetic trees inferred from mitochondrial genome (mitogenome) DNA sequences with nuclear data was investigated. The website Web of Science™ (Clarivate Analytics) was employed and the following search criteria: Topic [mito* genome] AND phylogeny NOT method$. Search restrictions were kept to a minimum to prevent unwanted exclusions leading to possible bias. The search was performed on November 23 of 2016 and yielded a total of 2,760 matches. The papers were ordered from newest to oldest to reflect the most recent trends. From these 2,760 published studies, the first 100 published studies which included mitogenome DNA sequences and employed those to estimate a mitochondrial phylogeny were assessed in further detail. The assessed 100 published studies that met these selection criteria are listed below.

**Results**

Among the 100 published studies assessed in more detail, only 14 completed the mitogenome DNA sequence-based phylogeny with data from nuclear loci. Four studies estimated a mitogenome DNA sequence-based phylogeny in a single species and 96 studies included multiple species even though the focus of the study was on a single species.

**Table S4. Manuscripts that include mitochondrial genome (mitogenome) data to reconstruct phylogenies and the use of complemented nuclear data**

| Web of Science study | Reference | Study assessed number | Mitogenome data | Nuclear data | Single species |
|---|---|---|---|---|---|
| 1 | (Aguado *et al.*, 2016) | 1 | YES (1) | | |
| 4 | (Li *et al.*, 2016b) | 2 | YES (1) | YES | |

| | | | | |
|---|---|---|---|---|
| 5 | (Soares *et al.*, 2016) | 3 | YES (1) | |
| 6 | (Song *et al.*, 2016c) | 4 | YES (1,3) | |
| 7 | (Liu *et al.*, 2016b) | 5 | YES (1,3) | |
| 8 | (Boo *et al.*, 2016) | 6 | YES (1) | |
| 12 | (Wu *et al.*, 2016b) | 7 | YES (1) | |
| 13 | (Zarza *et al.*, 2016) | 8 | YES (1) | YES |
| 14 | (Huang *et al.*, 2016) | 9 | YES (1,2) | |
| 16 | (Yong *et al.*, 2016c) | 10 | YES (1) | |
| 17 | (Sun *et al.*, 2016a) | 11 | YES (1) | |
| 18 | (Fortes *et al.*, 2016) | 12 | YES (4) | |
| 22 | (Capel *et al.*, 2016) | 13 | YES (1) | |
| 23 | (Wang *et al.*, 2016e) | 14 | YES (1) | |
| 24 | (Song *et al.*, 2016a) | 15 | YES (1) | |
| 25 | (Jiang *et al.*, 2016b) | 16 | YES (1) | |
| 26 | (Ma *et al.*, 2016a) | 17 | YES (1) | YES |
| 27 | (Song *et al.*, 2016b) | 18 | YES (1) | |
| 28 | (Wassermann *et al.*, 2016) | 19 | YES (1) | |
| 37 | (Mitchell *et al.*, 2016b) | 20 | YES (1) | |
| 38 | (Yong *et al.*, 2016a) | 21 | YES (1) | |
| 41 | (Chen *et al.*, 2016) | 22 | YES (1,3) | |
| 42 | (Romero *et al.*, 2016) | 23 | YES (1) | |
| 43 | (Ma *et al.*, 2016b) | 24 | YES (1,3) | |
| 47 | (Plazzi *et al.*, 2016) | 25 | YES (1) | |
| 48 | (Brabec *et al.*, 2016) | 26 | YES (1) | YES |
| 51 | (Nakajima *et al.*, 2016) | 27 | YES (1,3) | |

| | | | | | |
|---|---|---|---|---|---|
| 52 | (Shan *et al.*, 2016) | 28 | YES (1) | | |
| 55 | (Amaral *et al.*, 2016) | 29 | YES (1) | | |
| 58 | (Wang *et al.*, 2016d) | 30 | YES (1) | | |
| 59 | (Cheng *et al.*, 2016b) | 31 | YES (1,3) | | |
| 64 | (Cui *et al.*, 2016) | 32 | YES (1) | YES | |
| 65 | (Bellot *et al.*, 2016) | 33 | YES (1) | YES | |
| 66 | (Mitchell *et al.*, 2016a) | 34 | YES (4) | | |
| 67 | (Gao *et al.*, 2016) | 35 | YES (1,3) | | |
| 68 | (Zhou *et al.*, 2016) | 36 | YES (1) | YES | |
| 69 | (Shenkar *et al.*, 2016) | 37 | YES (1) | | |
| 71 | (Wang *et al.*, 2016b) | 38 | YES (1,3) | | |
| 79 | (Yong *et al.*, 2016d) | 39 | YES (1) | | |
| 82 | (Montiel *et al.*, 2016) | 40 | YES (2) | | YES |
| 83 | (Jiang *et al.*, 2016a) | 41 | YES (1) | | |
| 84 | (Zhang *et al.*, 2016a) | 42 | YES (1) | | |
| 89 | (Harrisson *et al.*, 2016) | 43 | YES (1) | | |
| 90 | (Wang *et al.*, 2016a) | 44 | YES (1,3) | | |
| 91 | (Briscoe *et al.*, 2016) | 45 | YES (1,3) | YES (6) | |
| 92 | (Wu *et al.*, 2016a) | 46 | YES (1,3) | | |
| 95 | (Oceguera-Figueroa *et al.*, 2016) | 47 | YES (1,3) | | |
| 96 | (Bocak *et al.*, 2016) | 48 | YES (1) | YES | |
| 97 | Ulfah *et al.* (2016) | 49 | YES (1) | YES | |
| 101 | (Lavikainen *et al.*, 2016) | 50 | YES (1) | YES (6) | |
| 103 | (Nadimi *et al.*, 2016) | 51 | YES (1) | | |

| | | | | | |
|---|---|---|---|---|---|
| 105 | (Sun *et al.*, 2016b) | 52 | YES (1) | | |
| 106 | (Kim *et al.*, 2016) | 53 | YES (1,3) | | |
| 109 | (Ramirez-Rios *et al.*, 2016) | 54 | YES (1) | | |
| 110 | (Hawkins *et al.*, 2016) | 55 | YES (2) | YES | |
| 112 | (Hirase *et al.*, 2016) | 56 | YES (2) | | YES |
| 113 | (Patra *et al.*, 2016) | 57 | YES (1) | | |
| 120 | (Bourguignon *et al.*, 2016) | 58 | YES (1) | | |
| 121 | (Yang *et al.*, 2016) | 59 | YES (5) | | |
| 123 | (Du *et al.*, 2016) | 60 | YES (1) | | |
| 141 | (Dodt *et al.*, 2016) | 61 | YES (1) | | |
| 150 | (Slater *et al.*, 2016) | 62 | YES (1) | | |
| 151 | (Cong *et al.*, 2016) | 63 | YES (1) | YES | |
| 154 | (Vyas *et al.*, 2016) | 64 | YES (2) | | YES |
| 155 | (Gibb *et al.*, 2016) | 65 | YES (1) | | |
| 156 | (Cheng *et al.*, 2016a) | 66 | YES (1,3) | | |
| 157 | (Uribe *et al.*, 2016) | 67 | YES (1) | | |
| 158 | (Shen *et al.*, 2016) | 68 | YES (1,3) | | |
| 161 | (Song *et al.*, 2016e) | 69 | YES (1) | | |
| 162 | (Uliano-Silva *et al.*, 2016) | 70 | YES (2) | | |
| 163 | (Rouse *et al.*, 2016) | 71 | YES (1,3) | YES | |
| 164 | (Yong *et al.*, 2016b) | 72 | YES (1) | | |
| 165 | (Diaz-Jaimes *et al.*, 2016) | 73 | YES (1) | | |
| 166 | (Zhao *et al.*, 2016) | 74 | YES (1) | | |
| 168 | (Liu *et al.*, 2016a) | 75 | YES (1,3) | | |
| 169 | (Zhang *et al.*, 2016b) | 76 | YES (1,3) | | |

| | | | | | |
|---|---|---|---|---|---|
| 171 | (Ning *et al.*, 2016) | 77 | YES (2) | | YES |
| 174 | (Zhang *et al.*, 2016c) | 78 | YES (1,3) | | |
| 178 | (Hou *et al.*, 2016) | 79 | YES (1) | | |
| 189 | (Botero-Castro *et al.*, 2016) | 80 | YES (1) | | |
| 196 | (Dray *et al.*, 2016b) | 81 | YES (1) | | |
| 197 | (Dray *et al.*, 2016a) | 82 | YES (1) | | |
| 198 | (Minton *et al.*, 2016) | 83 | YES (1) | | |
| 199 | (Huang & Tu, 2016) | 84 | YES (1) | | |
| 200 | (Li *et al.*, 2016a) | 85 | YES (1,5) | | |
| 201 | (Song *et al.*, 2016d) | 86 | YES (1) | | |
| 207 | (Ma *et al.*, 2016c) | 87 | YES (1) | | |
| 209 | (Balakirev *et al.*, 2016) | 88 | YES (1) | | |
| 216 | (Zang *et al.*, 2016) | 89 | YES (2) | | |
| 224 | (Guo *et al.*, 2016) | 90 | YES (1) | | |
| 229 | (Quezada-Romegialli *et al.*, 2016) | 91 | YES (1) | | |
| 230 | (Hu *et al.*, 2016) | 92 | YES (2) | | |
| 231 | (Jung *et al.*, 2016) | 93 | YES (1) | | |
| 232 | (Jeon *et al.*, 2016) | 94 | YES (1,3) | | |
| 233 | (DeHart *et al.*, 2016) | 95 | YES (1) | | |
| 235 | (Ni *et al.*, 2016) | 96 | YES (1) | | |
| 236 | (Choi *et al.*, 2016) | 97 | YES (2) | | |
| 238 | (Wang *et al.*, 2016c) | 98 | YES (1) | | |
| 239 | (Kumar *et al.*, 2016) | 99 | YES (1) | | |
| 240 | (Popovic *et al.*, 2016) | 100 | YES (2) | | |

Notes: Web of Science study: published study number based on the Web of Science search. Study assessed number: published study number among the 100 assessed studies. (1) Mitogenomic DNA sequences was used to conduct analyses based upon protein-coding regions and/or rRNA and tRNA regions. (2) Studies which based the estimation upon the complete mitogenomic DNA sequence to estimate phylogenies; or studies which failed to clearly specify which mitogenomic regions were employed. (3) Protein-coding regions were used to determine the amino acid sequence and subsequently employed to estimate the final phylogeny. (4) Studies reporting employing “a nearly complete mitogenome”. (5) Studies which were inaccessible and judged based on the contents in the abstract. (6) Ribosomal DNA was specified as the nuclear marker.

**References**

Aguado, M. T., Richter, S., Sontowski, R., Golombek, A., Struck, T. H., & Bleidorn, C. (2016). Syllidae mitochondrial gene order is unusually variable for Annelida. [Article]. *Gene, 594*(1), 89-96. doi: 10.1016/j.gene.2016.08.050

Amaral, D. T., Mitani, Y., Ohmiya, Y., & Viviani, V. R. (2016). Organization and comparative analysis of the mitochondrial genomes of bioluminescent Elateroidea (Coleoptera: Polyphaga). [Article]. *Gene, 586*(2), 254-262. doi: 10.1016/j.gene.2016.04.009

Árnason, U., Gullberg, A., & Widegren, B. (1991). The complete nucleotide sequence of the mitochondrial DNA of the fin whale, *Balaenoptera physalus*. *Journal of Molecular Evolution, 33*(6), 556-568. doi: 10.1007/bf02102808

Balakirev, E. S., Parensky, V. A., & Ayala, F. J. (2016). Complete mitochondrial genomes of the anadromous and resident forms of the lamprey Lethenteron camtschaticum. [Article]. *Mitochondrial DNA, 27*(3), 1730-1731. doi: 10.3109/19401736.2014.961143

Bellot, S., Cusimano, N., Luo, S. X., Sun, G. L., Zarre, S., Groger, A., . . . Renner, S. S. (2016). Assembled Plastid and Mitochondrial Genomes, as well as Nuclear Genes, Place the Parasite Family Cynomoriaceae in the Saxifragales. [Article]. *Genome Biology and Evolution, 8*(7), 2214-2230. doi: 10.1093/gbe/evw147

Bérubé, M., Aguilar, A., Dendanto, D., Larsen, F., Notarbartolo Di Sciara, G., Sears, R., . . . Palsbøll, P. J. (1998). Population genetic structure of North Atlantic, Mediterranean Sea and Sea of Cortez fin whales, *Balaenoptera physalus* (Linnaeus 1758): analysis of mitochondrial and nuclear loci. *Molecular Ecology, 7*(5), 585-599. doi: 10.1046/j.1365-294x.1998.00359.x

Bérubé, M., Jørgensen, H., McEwing, R., & Palsbøll, P. J. (2000). Polymorphic di-nucleotide microsatellite loci isolated from the humpback whale, *Megaptera novaeangliae*. *Molecular Ecology, 9*(12), 2181-2183. doi: 10.1046/j.1365-294X.2000.105315.x

Bérubé, M., Rew, M., Skaug, H., Jørgensen, H., Robbins, J., Best, P., . . . Palsbøll, P. (2005). Polymorphic microsatellite loci isolated from humpback whale, *Megaptera novaeangliae* and fin whale, *Balaenoptera physalus*. *Conservation Genetics, 6*(4), 631-636. doi: 10.1007/s10592-005-9017-5

Bocak, L., Kundrata, R., Fernandez, C. A., & Vogler, A. P. (2016). The discovery of Iberobaeniidae (Coleoptera: Elateroidea): a new family of beetles from Spain, with immatures detected by environmental DNA sequencing. [Article]. *Proceedings of the Royal Society B-Biological Sciences, 283*(1830). doi: 10.1098/rspb.2015.2350

Boo, G. H., Hughey, J. R., Miller, K. A., & Boo, S. M. (2016). Mitogenomes from type specimens, a genotyping tool for morphologically simple species: ten genomes of agar-producing red algae. [Article]. *Scientific Reports, 6*. doi: 10.1038/srep35337

Botero-Castro, F., Delsuc, F., & Douzery, E. J. P. (2016). Thrice better than once: quality control guidelines to validate new mitogenomes. [Article]. *Mitochondrial DNA, 27*(1), 449-454. doi: 10.3109/19401736.2014.900666

Bourguignon, T., Lo, N., Sobotnik, J., Sillam-Dusses, D., Roisin, Y., & Evans, T. A. (2016). Oceanic dispersal, vicariance and human introduction shaped the modern distribution of the termites Reticulitermes, Heterotermes and Coptotermes. [Article]. *Proceedings of the Royal Society B-Biological Sciences, 283*(1827). doi: 10.1098/rspb.2016.0179

Brabec, J., Kuchta, R., Scholz, T., & Littlewood, D. T. J. (2016). Paralogues of nuclear ribosomal genes conceal phylogenetic signals within the invasive Asian fish tapeworm lineage: evidence from next generation sequencing data. [Article]. *International Journal for Parasitology, 46*(9), 555-562. doi: 10.1016/j.ijpara.2016.03.009

Briscoe, A. G., Bray, R. A., Brabec, J., & Littlewood, D. T. J. (2016). The mitochondrial genome and ribosomal operon of Brachycladium goliath (Digenea: Brachycladiidae) recovered from a stranded minke whale. [Article]. *Parasitology International, 65*(3), 271-275. doi: 10.1016/j.parint.2016.02.004

Capel, K. C. C., Migotto, A. E., Zilberberg, C., Lin, M. F., Forsman, Z., Miller, D. J., & Kitahara, M. V. (2016). Complete mitochondrial genome sequences of Atlantic representatives of the invasive Pacific coral species Tubastraea coccinea and T. tagusensis (Scleractinia, Dendrophylliidae): Implications for species identification. [Article]. *Gene, 590*(2), 270-277. doi: 10.1016/j.gene.2016.05.034

Chen, Z. T., Mu, L. X., Wang, J. R., & Du, Y. Z. (2016). Complete Mitochondrial Genome of the Citrus Spiny Whitefly Aleurocanthus spiniferus (Quaintance) (Hemiptera: Aleyrodidae): Implications for the Phylogeny of Whiteflies. [Article]. *PloS one, 11*(8). doi: 10.1371/journal.pone.0161385

Cheng, T., Liu, G. H., Song, H. Q., Lin, R. Q., & Zhu, X. Q. (2016a). The complete mitochondrial genome of the dwarf tapeworm Hymenolepis nana-a neglected zoonotic helminth. [Article]. *Parasitology Research, 115*(3), 1253-1262. doi: 10.1007/s00436-015-4862-8

Cheng, X. F., Zhang, L. P., Yu, D. N., Storey, K. B., & Zhang, J. Y. (2016b). The complete mitochondrial genomes of four cockroaches (Insecta: Blattodea) and phylogenetic analyses within cockroaches. [Article]. *Gene, 586*(1), 115-122. doi: 10.1016/j.gene.2016.03.057

Choi, H. Y., Kim, T. W., & Kim, S. (2016). The complete mitochondrial genome of the Pacific needlefish Strongylura anastomella (Belonidae, Beloniformes) from Korea. [Article]. *Mitochondrial DNA, 27*(4), 2479-2480. doi: 10.3109/19401736.2015.1033707

Cong, Q., Shen, J. H., Warren, A. D., Borek, D., Otwinowski, Z., & Grishin, N. V. (2016). Speciation in Cloudless Sulphurs Gleaned from Complete Genomes. [Article]. *Genome Biology and Evolution, 8*(3), 915-931. doi: 10.1093/gbe/evw045

Cui, Y. Y., Yan, C. C., Sun, T. L., Li, J., Yue, B. S., & Zhang, X. Y. (2016). Identification of CR1 retroposons in Arborophila rufipectus and their application to Phasianidae phylogeny. [Article]. *Molecular ecology resources, 16*(4), 1037-1049. doi: 10.1111/1755-0998.12514

DeHart, H. M., Yang, L., & Naylor, G. J. P. (2016). Mitogenomic sequence and phylogenetic placement of the Hortle's whipray Himantura hortlei (Elasmobranchii: Dasyatidae). [Article]. *Mitochondrial DNA, 27*(4), 2437-2439. doi: 10.3109/19401736.2015.1030632

Diaz-Jaimes, P., Bayona-Vasquez, N. J., Adams, D. H., & Uribe-Alcocer, M. (2016). Complete mitochondrial DNA genome of bonnethead shark, Sphyrna tiburo, and phylogenetic relationships among main superorders of modern elasmobranchs. [Article]. *Meta Gene, 7*, 48-55. doi: 10.1016/j.mgene.2015.11.005

Dodt, W. G., McComish, B. J., Nilsson, M. A., Gibb, G. C., Penny, D., & Phillips, M. J. (2016). The complete mitochondrial genome of the eastern grey kangaroo (Macropus giganteus). [Article]. *Mitochondrial DNA, 27*(2), 1366-1367. doi: 10.3109/19401736.2014.947583

Dray, L., Neuhof, M., Diamant, A., & Huchon, D. (2016a). The complete mitochondrial genome of the devil firefish Pterois miles (Bennett, 1828) (Scorpaenidae). [Article]. *Mitochondrial DNA, 27*(1), 783-784. doi: 10.3109/19401736.2014.945565

Dray, L., Neuhof, M., Diamant, A., & Huchon, D. (2016b). The complete mitochondrial genome of the gilthead seabream Sparus aurata L. (Sparidae). [Article]. *Mitochondrial DNA, 27*(1), 781-782. doi: 10.3109/19401736.2014.928861

Du, C., He, S. L., Song, X. H., Liao, Q., Zhang, X. Y., & Yue, B. S. (2016). The complete mitochondrial genome of Epicauta chinensis (Coleoptera: Meloidae) and phylogenetic analysis among Coleopteran insects. [Article]. *Gene, 578*(2), 274-280. doi: 10.1016/j.gene.2015.12.036

Fortes, G. G., Grandal-d'Anglade, A., Kolbe, B., Fernandes, D., Meleg, I. N., Garcia-Vazquez, A., . . . Barlow, A. (2016). Ancient DNA reveals differences in behaviour and sociality between brown bears and extinct cave bears. [Article]. *Molecular Ecology, 25*(19), 4907-4918. doi: 10.1111/mec.13800

Gao, D. Z., Liu, G. H., Song, H. Q., Wang, G. L., Wang, C. R., & Zhu, X. Q. (2016). The complete mitochondrial genome of Gasterophilus intestinalis, the first representative of the family

Gasterophilidae. [Article]. *Parasitology Research, 115*(7), 2573-2579. doi: 10.1007/s00436-016-5002-9

Gibb, G. C., Condamine, F. L., Kuch, M., Enk, J., Moraes-Barros, N., Superina, M., . . . Delsuc, F. (2016). Shotgun Mitogenomics Provides a Reference Phylogenetic Framework and Timescale for Living Xenarthrans. [Article]. *Mol Biol Evol, 33*(3), 621-642. doi: 10.1093/molbev/msv250

Guo, Y. S., Bai, Q., Yan, T., Wang, Z. D., & Liu, C. W. (2016). Mitogenomes of genus Pristipomoides, Lutjanus and Pterocaesio confirm Caesionidae nests in Lutjanidae. [Article]. *Mitochondrial DNA, 27*(3), 2198-2199. doi: 10.3109/19401736.2014.982624

Harrisson, K., Pavlova, A., Gan, H. M., Lee, Y. P., Austin, C. M., & Sunnucks, P. (2016). Pleistocene divergence across a mountain range and the influence of selection on mitogenome evolution in threatened Australian freshwater cod species. [Article]. *Heredity, 116*(6), 506-515. doi: 10.1038/hdy.2016.8

Hawkins, M. T. R., Leonard, J. A., Helgen, K. M., McDonough, M. M., Rockwood, L. L., & Maldonado, J. E. (2016). Evolutionary history of endemic Sulawesi squirrels constructed from UCEs and mitogenomes sequenced from museum specimens. [Article]. *BMC evolutionary biology, 16*. doi: 10.1186/s12862-016-0650-z

Hirase, S., Takeshima, H., Nishida, M., & Iwasaki, W. (2016). Parallel Mitogenome Sequencing Alleviates Random Rooting Effect in Phylogeography. [Article]. *Genome Biology and Evolution, 8*(4), 1267-1278. doi: 10.1093/gbe/evw063

Hou, L. X., Ying, S., Yang, X. W., Yu, Z., Li, H. M., & Qin, X. M. (2016). The complete mitochondrial genome of Papilio bianor (Lepidoptera: Papilionidae), and its phylogenetic position within Papilionidae. [Article]. *Mitochondrial DNA, 27*(1), 102-103. doi: 10.3109/19401736.2013.873923

Hu, X. X., Lu, J., He, S. Y., Li, X. F., Liu, H. J., & Han, L. J. (2016). The complete mitochondrial genome of Astatotilapia burtoni. [Article]. *Mitochondrial DNA, 27*(4), 2379-2380. doi: 10.3109/19401736.2015.1028039

Huang, S. P., Chen, I. S., Jang-Liaw, N. H., Shao, K. T., & Yung, M. M. N. (2016). Complete Mitochondrial Reveals a New Phylogenetic Perspective on the Brackish Water Goby Mugilogobius Group (Teleostei: Gobiidae: Gobionellinae). [Article]. *Zoological Science, 33*(5), 566-574. doi: 10.2108/zs150154

Huang, Z. H., & Tu, F. Y. (2016). Mitogenome of Fejervarya multistriata: a novel gene arrangement and its evolutionary implications. [Article]. *Genetics and Molecular Research, 15*(3). doi: 10.4238/gmr.15038302

Jeon, M. G., Kim, J. Y., & Park, Y. C. (2016). Phylogenetic analysis of the complete mitochondrial genome of the Korean field mouse Apodemus peninsulae (Rodentia, Murinae) from China. [Article]. *Mitochondrial DNA, 27*(4), 2408-2409. doi: 10.3109/19401736.2015.1030618

Jiang, P., Li, H., Song, F., Cai, Y., Wang, J. Y., Liu, J. P., & Cai, W. Z. (2016a). Duplication and Remolding of tRNA Genes in the Mitochondrial Genome of Reduvius tenebrosus (Hemiptera: Reduviidae). [Article]. *International Journal of Molecular Sciences, 17*(6). doi: 10.3390/ijms17060951

Jiang, Y. L., Yang, F., Yang, D., & Liu, X. Y. (2016b). Complete mitochondrial genome of a Neotropical dobsonfly Chloronia mirifica Navas, 1925 (Megaloptera: Corydalidae), with phylogenetic implications for the genus Chloronia Banks, 1908. [Article]. *Zootaxa, 4162*(1), 46-60. doi: 10.11646/zootaxa.4162.1.2

Jung, G., Kim, C. G., & Lee, Y. H. (2016). Complete mitochondrial genome sequence of Heliocidaris crassispina (Camarodonta, Echinometridae). [Article]. *Mitochondrial DNA, 27*(4), 2393-2394. doi: 10.3109/19401736.2015.1028046

Kim, T., Kim, J., Nadler, S. A., & Park, J. K. (2016). The complete mitochondrial genome of Koerneria sudhausi (Diplogasteromorpha: Nematoda) supports monophyly of Diplogasteromorpha within Rhabditomorpha. [Article]. *Current Genetics, 62*(2), 391-403. doi: 10.1007/s00294-015-0536-4

Kumar, R., Goel, C., Sahoo, P. K., Singh, A. K., & Barat, A. (2016). Complete mitochondrial genome organization of Tor tor (Hamilton, 1822). [Article]. *Mitochondrial DNA, 27*(4), 2541-2542. doi: 10.3109/19401736.2015.1038795

Lavikainen, A., Iwaki, T., Haukisalmi, V., Konyaev, S. V., Casiraghi, M., Dokuchaev, N. E., . . . Nakao, M. (2016). Reappraisal of Hydatigera taeniaeformis (Batsch, 1786) (Cestoda: Taeniidae) sensu lato with description of Hydatigera kamiyai n. sp. [Article]. *International Journal for Parasitology, 46*(5-6), 361-374. doi: 10.1016/j.ijpara.2016.01.009

Li, C. L., Du, X. Y., Gao, J., Wang, C., Guo, H. G., Dai, F. W., . . . Chen, Z. W. (2016a). Phylogenetic analysis of the Mongolian gerbil (Meriones unguiculatus) from China based on mitochondrial genome. [Article]. *Genetics and Molecular Research, 15*(3). doi: 10.4238/gmr.15037703

Li, X. J., Lin, L. L., Cui, A. M., Bai, J., Wang, X. Y., Xin, C., . . . Lei, F. M. (2016b). Taxonomic status and phylogenetic relationship of tits based on mitogenomes and nuclear segments. [Article]. *Mol Phylogenet Evol, 104*, 14-20. doi: 10.1016/j.ympev.2016.07.022

Liu, Q. N., Chai, X. Y., Bian, D. D., Ge, B. M., Zhou, C. L., & Tang, B. P. (2016a). The complete mitochondrial genome of fall armyworm Spodoptera frugiperda (Lepidoptera: Noctuidae). [Article]. *Genes & Genomics, 38*(2), 205-216. doi: 10.1007/s13258-015-0346-6

Liu, Q. N., Xin, Z. Z., Bian, D. D., Chai, X. Y., Zhou, C. L., & Tang, B. P. (2016b). The first complete mitochondrial genome for the subfamily Limacodidae and implications for the higher phylogeny of Lepidoptera. [Article]. *Scientific Reports, 6*. doi: 10.1038/srep35878

Ma, H. F., Zheng, X. X., Peng, M. H., Bian, H. X., Chen, M. M., Liu, Y. Q., . . . Qin, L. (2016a). Complete mitochondrial genome of the meadow moth, Loxostege sticticalis (Lepidoptera: Pyraloidea: Crambidae), compared to other Pyraloidea moths. [Article]. *Journal of Asia-Pacific Entomology, 19*(3), 697-706. doi: 10.1016/j.aspen.2016.05.011

Ma, J., He, J. J., Liu, G. H., Leontovyc, R., Kasny, M., & Zhu, X. Q. (2016b). Complete mitochondrial genome of the giant liver fluke Fascioloides magna (Digenea: Fasciolidae) and its comparison with selected trematodes. [Article]. *Parasites & Vectors, 9*. doi: 10.1186/s13071-016-1699-7

Ma, Q. Z., Wu, B., Li, J. X., & Song, Z. B. (2016c). Complete mitochondrial genome of Ptychobarbus kaznakovi (Teleostei: Cypriniformes: Cyprinidae), and repetitive sequences in the D-loop. [Article]. *Mitochondrial DNA, 27*(3), 1683-1684. doi: 10.3109/19401736.2014.958727

Minton, R. L., Cruz, M. A. M., Farman, M. L., & Perez, K. E. (2016). Two complete mitochondrial genomes from Praticolella mexicana Perez, 2011 (Polygyridae) and gene order evolution in Helicoidea (Mollusca, Gastropoda). [Article]. *Zookeys*(626), 137-154. doi: 10.3897/zookeys.626.9633

Mitchell, K. J., Scanferla, A., Soibelzon, E., Bonini, R., Ochoa, J., & Cooper, A. (2016a). Ancient DNA from the extinct South American giant glyptodont Doedicurus sp (Xenarthra: Glyptodontidae) reveals that glyptodonts evolved from Eocene armadillos. [Article]. *Molecular Ecology, 25*(14), 3499-3508. doi: 10.1111/mec.13695

Mitchell, K. J., Wood, J. R., Llamas, B., McLenachan, P. A., Kardailsky, O., Scofield, R. P., . . . Cooper, A. (2016b). Ancient mitochondrial genomes clarify the evolutionary history of New Zealand's enigmatic acanthisittid wrens. [Article]. *Mol Phylogenet Evol, 102*, 295-304. doi: 10.1016/j.ympev.2016.05.038

Montiel, A. L., Hail, D., Macias-Velasco, J. F., Powell, C. M., & Bextine, B. R. (2016). The Mitochondrial Genome of the Potato Psyllid, Bactericera cockerelli Sulc., and Differences among Potato Psyllid Populations of the United States. [Article]. *Southwestern Entomologist, 41*(2), 347-360.

Nadimi, M., Daubois, L., & Hijri, M. (2016). Mitochondrial comparative genomics and phylogenetic signal assessment of mtDNA among arbuscular mycorrhizal fungi. [Article]. *Mol Phylogenet Evol, 98*, 74-83. doi: 10.1016/j.ympev.2016.01.009

Nakajima, Y., Shinzato, C., Khalturina, M., Nakamura, M., Watanabe, H., Satoh, N., & Mitarai, S. (2016). The mitochondrial genome sequence of a deep-sea, hydrothermal vent limpet,

Lepetodrilus nux, presents a novel vetigastropod gene arrangement. [Article]. *Marine Genomics, 28*, 121-126. doi: 10.1016/j.margen.2016.04.005

Ni, N. N., Yu, D. N., Storey, K. B., Zheng, R. Q., & Zhang, J. Y. (2016). The complete mitochondrial genome of Lithobates sylvaticus (Anura: Ranidae). [Article]. *Mitochondrial DNA, 27*(4), 2460-2461. doi: 10.3109/19401736.2015.1033697

Ning, C., Gao, S. Z., Deng, B. P., Zheng, H. X., Wei, D., Lv, H. Z., . . . Cui, Y. Q. (2016). Ancient mitochondrial genome reveals trace of prehistoric migration in the east Pamir by pastoralists. [Article]. *Journal of Human Genetics, 61*(2), 103-108. doi: 10.1038/jhg.2015.128

Oceguera-Figueroa, A., Manzano-Marin, A., Kvist, S., Moya, A., Siddall, M. E., & Latorre, A. (2016). Comparative Mitogenomics of Leeches (Annelida: Clitellata): Genome Conservation and Placobdella-Specific trnD Gene Duplication. [Article]. *PloS one, 11*(5). doi: 10.1371/journal.pone.0155441

Palsbøll, P. J., Bérubé, M., Larsen, A. H., & Jørgensen, H. (1997). Primers for the amplification of tri- and tetramer microsatellite loci in baleen whales. *Molecular Ecology, 6*(9), 893-895. doi: 10.1046/j.1365-294X.1997.d01-214.x

Patra, A. K., Kwon, Y. M., Kang, S. G., Fujiwara, Y., & Kim, S. J. (2016). The complete mitochondrial genome sequence of the tubeworm Lamellibrachia satsuma and structural conservation in the mitochondrial genome control regions of Order Sabellida. [Article]. *Marine Genomics, 26*, 63-71. doi: 10.1016/j.margen.2015.12.010

Plazzi, F., Puccio, G., & Passamonti, M. (2016). Comparative Large-Scale Mitogenomics Evidences Clade-Specific Evolutionary Trends in Mitochondrial DNAs of Bivalvia. [Article]. *Genome Biology and Evolution, 8*(8), 2544-2564. doi: 10.1093/gbe/evw187

Popovic, D., Baca, M., & Panagiotopoulou, H. (2016). Complete mitochondrial genome sequences of Atlantic sturgeon, Acipenser oxyrinchus oxyrinchus, Gulf sturgeon, A-o. desotoi and European sturgeon A-sturio (Acipenseriformes: Acipenseridae) obtained through next generation sequencing. [Article]. *Mitochondrial DNA, 27*(4), 2549-2551. doi: 10.3109/19401736.2015.1038799

Quezada-Romegialli, C., Veliz, D., Docmac, F., & Harrod, C. (2016). The complete mitochondrial genome of the rocky reef fish Cheilodactylus variegatus Valenciennes, 1833 (Teleostei: Cheilodactylidae). [Article]. *Mitochondrial DNA, 27*(4), 2359-2360. doi: 10.3109/19401736.2015.1025263

Ramirez-Rios, V., Franco-Sierra, N. D., Alvarez, J. C., Saldamando-Benjumea, C. I., & Villanueva-Mejia, D. F. (2016). Mitochondrial genome characterization of Tecia solanivora (Lepidoptera: Gelechiidae) and its phylogenetic relationship with other lepidopteran insects. [Article]. *Gene, 581*(2), 107-116. doi: 10.1016/j.gene.2016.01.031

Romero, P. E., Weigand, A. M., & Pfenninger, M. (2016). Positive selection on panpulmonate mitogenomes provide new clues on adaptations to terrestrial life. [Article]. *BMC evolutionary biology, 16*. doi: 10.1186/s12862-016-0735-8

Rouse, G. W., Wilson, N. G., Carvajal, J. I., & Vrijenhoek, R. C. (2016). New deep-sea species of Xenoturbella and the position of Xenacoelomorpha. [Article]. *Nature, 530*(7588), 94-+. doi: 10.1038/nature16545

Shan, B. B., Song, N., Han, Z. Q., Wang, J., Gao, T. X., & Yokogawa, K. (2016). Complete mitochondrial genomes of three sea basses Lateolabrax (Perciformes, Lateolabracidae) species: Genome description and phylogenetic considerations. [Article]. *Biochemical Systematics and Ecology, 67*, 44-52. doi: 10.1016/j.bse.2016.04.007

Shen, X., Sun, S., Zhao, F. Q., Zhang, G. T., Tian, M., Tsang, L. M., . . . Chu, K. H. (2016). Phylomitogenomic analyses strongly support the sister relationship of the Chaetognatha and Protostomia. [Article]. *Zoologica Scripta, 45*(2), 187-199. doi: 10.1111/zsc.12140

Shenkar, N., Koplovitz, G., Dray, L., Gissi, C., & Huchon, D. (2016). Back to solitude: Solving the phylogenetic position of the Diazonidae using molecular and developmental characters. [Article]. *Mol Phylogenet Evol, 100*, 51-56. doi: 10.1016/j.ympev.2016.04.001

Slater, G. J., Cui, P., Forasiepi, A. M., Lenz, D., Tsangaras, K., Voirin, B., . . . Greenwood, A. D. (2016). Evolutionary Relationships among Extinct and Extant Sloths: The Evidence of Mitogenomes and Retroviruses. [Article]. *Genome Biology and Evolution, 8*(3), 607-621. doi: 10.1093/gbe/evw023

Soares, A. E. R., Novak, B. J., Haile, J., Heupink, T. H., Fjeldsa, J., Gilbert, M. T. P., . . . Shapiro, B. (2016). Complete mitochondrial genomes of living and extinct pigeons revise the timing of the columbiform radiation. [Article]. *BMC evolutionary biology, 16*. doi: 10.1186/s12862-016-0800-3

Song, N., An, S., Yin, X., Cai, W., & Li, H. (2016a). Application of RNA-seq for mitogenome reconstruction, and reconsideration of long-branch artifacts in Hemiptera phylogeny. *Scientific Reports, 6*.

Song, N., An, S. H., Yin, X. M., Zhao, T., & Wang, X. Y. (2016b). Insufficient resolving power of mitogenome data in deciphering deep phylogeny of Holometabola. [Article]. *Journal of Systematics and Evolution, 54*(5), 545-559. doi: 10.1111/jse.12214

Song, N., Li, H., Song, F., & Cai, W. Z. (2016c). Molecular phylogeny of Polyneoptera (Insecta) inferred from expanded mitogenomic data. [Article]. *Scientific Reports, 6*. doi: 10.1038/srep36175

Song, S. L., Yong, H. S., Lim, P. E., Ng, P. K., & Phang, S. M. (2016d). Complete mitochondrial genome, genetic diversity and molecular phylogeny of Gracilaria salicornia (Rhodophyta: Gracilariaceae). [Article]. *Phycologia, 55*(4), 371-377. doi: 10.2216/15-128.1

Song, S. N., Tang, P., Wei, S. J., & Chen, X. X. (2016e). Comparative and phylogenetic analysis of the mitochondrial genomes in basal hymenopterans. [Article]. *Scientific Reports, 6*. doi: 10.1038/srep20972

Sun, M. M., Ma, J., Sugiyama, H., Ando, K., Li, W. W., Xu, Q. M., . . . Zhu, X. Q. (2016a). The complete mitochondrial genomes of Gnathostoma doloresi from China and Japan. [Article]. *Parasitology Research, 115*(10), 4013-4020. doi: 10.1007/s00436-016-5171-6

Sun, Y., Jiang, Q., Yang, C. H., Wang, X. G., Tian, F., Wang, Y. C., . . . Wang, C. F. (2016b). Characterization of complete mitochondrial genome of Dezhou donkey (Equus asinus) and evolutionary analysis. [Article]. *Current Genetics, 62*(2), 383-390. doi: 10.1007/s00294-015-0531-9

Ulfah, M., Kawahara-Miki, R., Farajalllah, A., Muladno, M., Dorshorst, B., Martin, A., & Kono, T. (2016). Genetic features of red and green junglefowls and relationship with Indonesian native chickens Sumatera and Kedu Hitam. [Article]. *Bmc Genomics, 17*. doi: 10.1186/s12864-016-2652-z

Uliano-Silva, M., Americo, J. A., Costa, I., Schomaker-Bastos, A., Rebelo, M. D., & Prosdocimi, F. (2016). The complete mitochondrial genome of the golden mussel Limnoperna fortunei and comparative mitogenomics of Mytilidae. [Article]. *Gene, 577*(2), 202-208. doi: 10.1016/j.gene.2015.11.043

Uribe, J. E., Kano, Y., Templado, J., & Zardoya, R. (2016). Mitogenomics of Vetigastropoda: insights into the evolution of pallial symmetry. *Zoologica Scripta, 45*(2), 145-159.

Valsecchi, E., & Amos, W. (1996). Microsatellite markers for the study of cetacean populations. *Molecular Ecology, 5*(1), 151-156. doi: 10.1111/j.1365-294X.1996.tb00301.x

Vyas, D. N., Kitchen, A., Miro-Herrans, A. T., Pearson, L. N., Al-Meeri, A., & Mulligan, C. J. (2016). Bayesian analyses of Yemeni mitochondrial genomes suggest multiple migration events with Africa and Western Eurasia. [Article]. *American Journal of Physical Anthropology, 159*(3), 382-393. doi: 10.1002/ajpa.22890

Wang, B. J., Gu, X. B., Yang, G. Y., Wang, T., Lai, W. M., Zhong, Z. J., & Liu, G. H. (2016a). Mitochondrial genomes of Heterakis gallinae and Heterakis beramporia support that they belong to the infraorder Ascaridomorpha. [Article]. *Infection Genetics and Evolution, 40*, 228-235. doi: 10.1016/j.meegid.2016.03.012

Wang, C. R., Lou, Y., Gao, J. F., Qiu, J. H., Zhang, Y., Gao, Y., & Chang, Q. C. (2016b). Comparative analyses of the complete mitochondrial genomes of the two murine pinworms Aspiculuris tetraptera and Syphacia obvelata. [Article]. *Gene, 585*(1), 71-75. doi: 10.1016/j.gene.2016.03.037

Wang, G. P., Min, Q., & Si, G. C. (2016c). The complete mitochondrial genome sequence of Pterophyllum scalare (Cichliformes: Cichlidae). [Article]. *Mitochondrial DNA, 27*(4), 2506-2507. doi: 10.3109/19401736.2015.1036251

Wang, Y., Shen, Y. J., Feng, C. G., Zhao, K., Song, Z. B., Zhang, Y. P., . . . He, S. P. (2016d). Mitogenomic perspectives on the origin of Tibetan loaches and their adaptation to high altitude. [Article]. *Scientific Reports, 6*. doi: 10.1038/srep29690

Wang, Z. L., Li, C., Fang, W. Y., & Yu, X. P. (2016e). Characterization of the complete mitogenomes of two Neoscona spiders (Araneae: Araneidae) and its phylogenetic implications. [Article]. *Gene, 590*(2), 298-306. doi: 10.1016/j.gene.2016.05.037

Wassermann, M., Woldeyes, D., Gerbi, B. M., Ebi, D., Zeyhle, E., Mackenstedt, U., . . . Romig, T. (2016). A novel zoonotic genotype related to Echinococcus granulosus sensu stricto from southern Ethiopia. [Article]. *International Journal for Parasitology, 46*(10), 663-668. doi: 10.1016/j.ijpara.2016.04.005

Wu, F. N., Cen, Y. J., Wallis, C. M., Trumble, J. T., Prager, S., Yokomi, R., . . . Liang, G. W. (2016a). The Complete Mitochondrial Genome Sequence of Bactericera cockerelli and Comparison with Three Other Psylloidea Species. [Article]. *PloS one, 11*(5). doi: 10.1371/journal.pone.0155318

Wu, Y. P., Zhao, J. L., Su, T. J., Luo, A. R., & Zhu, C. D. (2016b). The complete mitochondrial genome of Choristoneura longicellana (Lepidoptera: Tortricidae) and phylogenetic analysis of Lepidoptera. [Article]. *Gene, 591*(1), 161-176. doi: 10.1016/j.gene.2016.07.003

Yang, J., Ren, Q. L., Zhang, Q., & Huang, Y. (2016). Complete mitochondrial genomes of three crickets (Orthoptera: Gryllidae) and comparative analyses within Ensifera mitogenomes. [Article]. *Zootaxa, 4092*(4), 529-547.

Yong, H. S., Song, S. L., Eamsobhana, P., & Lim, P. E. (2016a). Complete mitochondrial genome of Angiostrongylus malaysiensis lungworm and molecular phylogeny of Metastrongyloid nematodes. [Article]. *Acta Tropica, 161*, 33-40. doi: 10.1016/j.actatropica.2016.05.002

Yong, H. S., Song, S. L., Lim, P. E., Eamsobhana, P., & Suana, I. W. (2016b). Complete Mitochondrial Genome of Three Bactrocera Fruit Flies of Subgenus Bactrocera (Diptera: Tephritidae) and Their Phylogenetic Implications. [Article]. *PloS one, 11*(2). doi: 10.1371/journal.pone.0148201

Yong, H. S., Song, S. L., Lim, P. E., Eamsobhana, P., & Suana, I. W. (2016c). Differentiating sibling species of Zeugodacus caudatus (Insecta: Tephritidae) by complete mitochondrial genome. [Article]. *Genetica, 144*(5), 513-521. doi: 10.1007/s10709-016-9919-9

Yong, H. S., Song, S. L., Lim, P. E., Eamsobhana, P., & Tan, J. (2016d). Complete mitochondrial genome and phylogeny of Microhyla butleri (Amphibia: Anura: Microhylidae). [Article]. *Biochemical Systematics and Ecology, 66*, 243-253. doi: 10.1016/j.bse.2016.04.004

Zang, X., Wang, X. J., Zhang, G. S., Wang, Y. Y., Ding, Y. D., & Yin, S. W. (2016). Complete mitochondrial genome and phylogenic analysis of Odontobutis yaluensis, Perciformes, Odontobutidae. [Article]. *Mitochondrial DNA, 27*(3), 1965-1967. doi: 10.3109/19401736.2014.971309

Zarza, E., Faircloth, B. C., Tsai, W. L. E., Bryson, R. W., Klicka, J., & McCormack, J. E. (2016). Hidden histories of gene flow in highland birds revealed with genomic markers. [Article]. *Molecular Ecology, 25*(20), 5144-5157. doi: 10.1111/mec.13813

Zhang, H. L., Liu, B. B., Wang, X. Y., Han, Z. P., Zhang, D. X., & Su, C. N. (2016a). Comparative Mitogenomic Analysis of Species Representing Six Subfamilies in the Family Tenebrionidae. [Article]. *International Journal of Molecular Sciences, 17*(6). doi: 10.3390/ijms17060841

Zhang, J., Chen, Z., Zhou, C. J., & Kong, X. H. (2016b). Molecular phylogeny of the subfamily Schizothoracinae (Teleostei: Cypriniformes: Cyprinidae) inferred from complete mitochondrial genomes. [Article]. *Biochemical Systematics and Ecology, 64*, 6-13. doi: 10.1016/j.bse.2015.11.004

Zhang, L. L., Sechi, P., Yuan, M. L., Jiang, J. B., Dong, Y., & Qiu, J. P. (2016c). Fifteen new earthworm mitogenomes shed new light on phylogeny within the Pheretima complex. [Article]. *Scientific Reports, 6*. doi: 10.1038/srep20096

Zhao, C., Zhang, H. H., Liu, G. S., Yang, X. F., & Zhang, J. (2016). The complete mitochondrial genome of the Tibetan fox (Vulpes ferrilata) and implications for the phylogeny of Canidae. [Article]. *Comptes Rendus Biologies, 339*(2), 68-77. doi: 10.1016/j.crvi.2015.11.005

Zhou, C. J., Wang, X. Z., Gan, X. N., Zhang, Y. P., Irwin, D. M., Mayden, R. L., & He, S. P. (2016). Diversification of Sisorid catfishes (Teleostei: Siluriformes) in relation to the orogeny of the Himalayan Plateau. [Article]. *Science Bulletin, 61*(13), 991-1002. doi: 10.1007/s11434-016-1104-0
